# Single-cell profiling guided combinatorial immunotherapy for fast-evolving CDK4/6 inhibitor resistant HER2-positive breast cancer

**DOI:** 10.1101/671198

**Authors:** Qingfei Wang, Ian H. Guldner, Samantha M. Golomb, Longhua Sun, Jack Harris, Xin Lu, Siyuan Zhang

**Author notes:** Correspondence and requests for materials should be addressed to Q.W. or to S.Z.

## Abstract

Development of acquired resistance to targeted cancer therapy is one of the most significant clinical challenges. Acquiring resistance under drug selection pressure is a result of evolutionary adaptation to a complex and dynamic tumor microenvironment (TME). New therapy regimens combining CDK4/6 inhibitor are under active investigation in clinical trials to treat HER2+ breast cancer patients. In parallel with clinical trial settings, in this study, we sought to prospectively model the tumor evolution in response to a targeted therapy regimen *in vivo* and identify a clinically actionable strategy to combat potential acquired resistance. Notably, despite a promising initial response, acquired resistance emerged rapidly to the anti-Her2/Neu antibody plus CDK4/6 inhibitor Palbociclib combination treatment. By leveraging high-throughput single-cell analyses of the evolving tumors over the course of treatments, we revealed a distinct immunosuppressive immature myeloid cell (IMC) population infiltrated in the resistant TME. Guided by single-cell transcriptome analysis, we demonstrated a combinatorial immunotherapy of IMC-targeting tyrosine kinase inhibitor cabozantinib and immune checkpoint blockades enhanced anti-tumor immunity, and overcame the resistance. Further, sequential combinatorial immunotherapy enabled a sustained control of the rapidly evolving CDK4/6 inhibitor-resistant tumors. Our study demonstrates a translational framework for treating rapidly evolving tumors through preclinical modeling and single-cell analyses. Our findings provide a rationale for an immediate clinical proposition of combinatorial immunotherapy for HER2+ breast cancer as a strategy to mitigate the emergence of resistance.

## Introduction

Precision medicine is a personalized disease treatment model that tailors the therapeutic regimens for individual patients by considering the genetic heterogeneity of disease ^1,2^. Targeted cancer therapy exemplifies the concept of precision medicine through rationally-designed targeted treatment towards a tumor-specific dysregulated genetic event ^3,4^. With superior clinical efficacy and less side effects, targeted cancer therapy has become one of the major pillars of modern cancer treatment ^5,6^. However, cancer is a consistently evolving multicellular ecosystem ^7^. Despite the initial clinical response to targeted therapies, drug-resistant tumors often emerge after prolonged treatments, which imposes a significant clinical challenge ^8,9^. Significant intratumoral heterogeneity of the tumor ecosystem, at both genetic and phenotypical level, is one of the primary culprits responsible for emergence of drug resistant tumors under the selection pressure of targeted therapy^10^. Majority of the research efforts on the resistance mechanisms have focused on such intratumoral heterogeneity of tumor cells, demonstrating that emergence of the drug-resistant phenotype is a result of selecting rare tumor cells with either pre-existing mutations (*de novo*) or newly acquired mutations (acquired) that confer resistance to specific targeted therapies ^11^. In addition to the tumor cell centric mechanisms, emerging evidence started to reveal that tumor microenvironment (TME) factors (e.g. biophysical/biochemical clues, stromal cells), collaboratively contribute to the evolving path of the tumor to seemingly inevitable resistance ^12^.

Modeling the dynamic nature of evolving drug resistance while capturing a holistic view of both tumor cells and the TME is essential for a systematic interrogation of resistance mechanisms and designing novel strategies to overcome resistance ^12–14^. Traditionally, exploring molecular underpinnings of drug resistance relies on either one-pathway-at-a-time approach using *in vitro* cell culture model or bulk DNA/RNA sequencing approaches comparing drug sensitive/responsive and resistant clinical tumor samples ^15,16^. However, the *in vitro* models cannot capture the interplay between evolving tumor cells and the microenvironment, and bulk sequencing has limited resolution in revealing tumor heterogeneity or identifying rare cellular events that confer phenotypical significance to drug resistance ^17^. Recent advances of single-cell analyses are revolutionizing the traditional paradigm of studying drug resistance by enabling a more holistic interrogation of tumor progression in response to drug treatments at an unprecedented single cell resolution ^18–20^. Single-cell sequencing approaches have effectively revealed intratumoral subclonal hierarchy at diagnosis ^21^, Darwinian clonal repopulation^19,22^, epigenetic reprogramming associated with resistant tumor cells ^23^ and dynamic changes of tumor-associated immune landscape ^18,24–27^. These pioneering studies based on single-cell analyses start to shed light on future single-cell analysis-based clinical management strategies for patients with relapsed resistant tumors ^28^.

Trastuzumab, widely known as Herceptin™, a humanized monoclonal antibody targeting the extracellular domain of human epidermal growth factor receptor-2 (HER2), is one of the most successful examples of targeted therapies for HER2-overexpressing breast cancer ^29^. Despite its significant initial therapeutic efficacy, both *de novo* and acquired resistance to trastuzumab have been observed in certain patients ^30,31^. Using *in vitro* trastuzumab-resistant cell line model, pre-clinical studies have mechanistically defined diverse intracellular signaling events conferring resistance, including but not limited to truncation of the HER2 receptor, dysregulating of PI3K/PTEN pathway, and engaging alternative survival pathways ^31^. Recently, using an inducible-HER2 transgenic mouse model, Goel et.al. revealed an enhanced cyclin D1-CDK4 dependent proliferation confers trastuzumab-resistance *in vivo* ^32^. Targeting cyclin D1-CDK4 acts synergistically with trastuzumab and, more intriguingly, elicits anti-tumor immune response ^33,34^. In light of such strong preclinical evidence and together with the recent accelerated FDA approval of CDK4/6 inhibitors for estrogen receptor (ER)–positive breast cancer patients (Palbociclib, Pfizer, FDA 2015, Ribociclib, Novartis, FDA 2017, and Abemaciclib, Lilly, FDA 2017), the new combinatorial regimen of CDK4/6 inhibitors plus trastuzumab is currently under active clinical investigation ^35,36^. Despite the clinical promise of CDK4/6 inhibitor containing regimen in treating HER2+ breast cancer, one can envision that the therapeutic resistance to anti-CDK4/6 treatment will ultimately emerge. Thus, in light with current clinical trials, we reasoned that prospectively modeling the tumor evolution in response to a trastuzumab plus CDK4/6 inhibitor regimen will provide valuable insight to the potential acquired resistance mechanisms. Preclinically, proactively exploring alternative therapeutic strategies that target emerging resistance mechanisms to prevent or inhibit resistance will have a direct translational impact on ongoing clinical trials and improve the therapeutic outcome.

In this study, we prospectively modeled *in vivo* acquired resistance to CDK4/6 inhibitor plus trastuzumab regimen using a transgenic mouse model in parallel with the current clinical trial scenario. We found that acquired resistance to the anti-Her2/Neu antibody plus Palbociclib combination (Ab+Pal) treatment emerged quickly after initial response. Through high-throughput single-cell RNA-seq and mass cytometry by time of flight (CyTOF) analyses of the evolving tumors over the course of treatment, including treatment naive, treatment responsive/residual disease and rapidly relapsed tumors, we revealed a distinct immunosuppressive immature myeloid cell (IMC) population infiltrated in the resistant TME. Next, guided by single-cell analyses, we evaluated the *in vivo* efficacy of using combinatorial immunotherapy by concomitantly targeting IMCs and enhancing T-cell activity. Further, our rationally designed sequential combinatorial regimens enabled a durable response and sustained control of the emergence of acquired resistance in rapidly evolving HER2-positive breast cancers.

## Results

### Rapid emergence of resistant tumors *in vivo* to anti-Her2/neu and CDK4/6 combinatorial targeted therapy with an increased antigen presentation and interferon signaling

To address the question whether long term Her2/neu and CDK4/6 inhibition in advanced HER2-positive breast cancer has a sustainable therapeutic effect, we employed the MMTV-neu202^Mul^ transgenic mouse bearing late-stage mammary tumor (volume > 500mm^3^) and examined their response to a continuous anti-Her2/neu antibody (Ab) plus CDK4/6 inhibitor Palbociclib (Pal) treatment. Two weeks of Ab+Pal treatment produced pronounced effects, leading to tumor regression with an average volume reduction of 52.74% (Fig. 1A) and significant suppression of tumor cell proliferation (Supplemental Fig. 1A). In contrast, control mice exhibited an average of 108.4% increase in tumor size over the same period, and Pal or Ab single treatment only showed a mild to moderate therapeutic effect (Fig. 1A and Supplemental Fig. 1A). Despite the initial significant efficacy of Ab+Pal combination and extended survival to doubled tumor volume (Supplemental Fig. 1B), shortly after tumor regression (2-4 weeks post-treatment), all combination treated tumors rebounded and eventually developed resistance under a long-term trial of Ab+Pal combination (Fig. 1B, Supplemental Fig. 1, C and D). The resistant tumors (Ab+Pal resistant, APR) exhibited higher proliferation as indicated by Ki67 staining compared to control tumor (Supplemental Fig. 1E), indicating a state of rapid growth of the resistant tumor even under the pressure of Ab+Pal treatment.

**Fig. 1.**
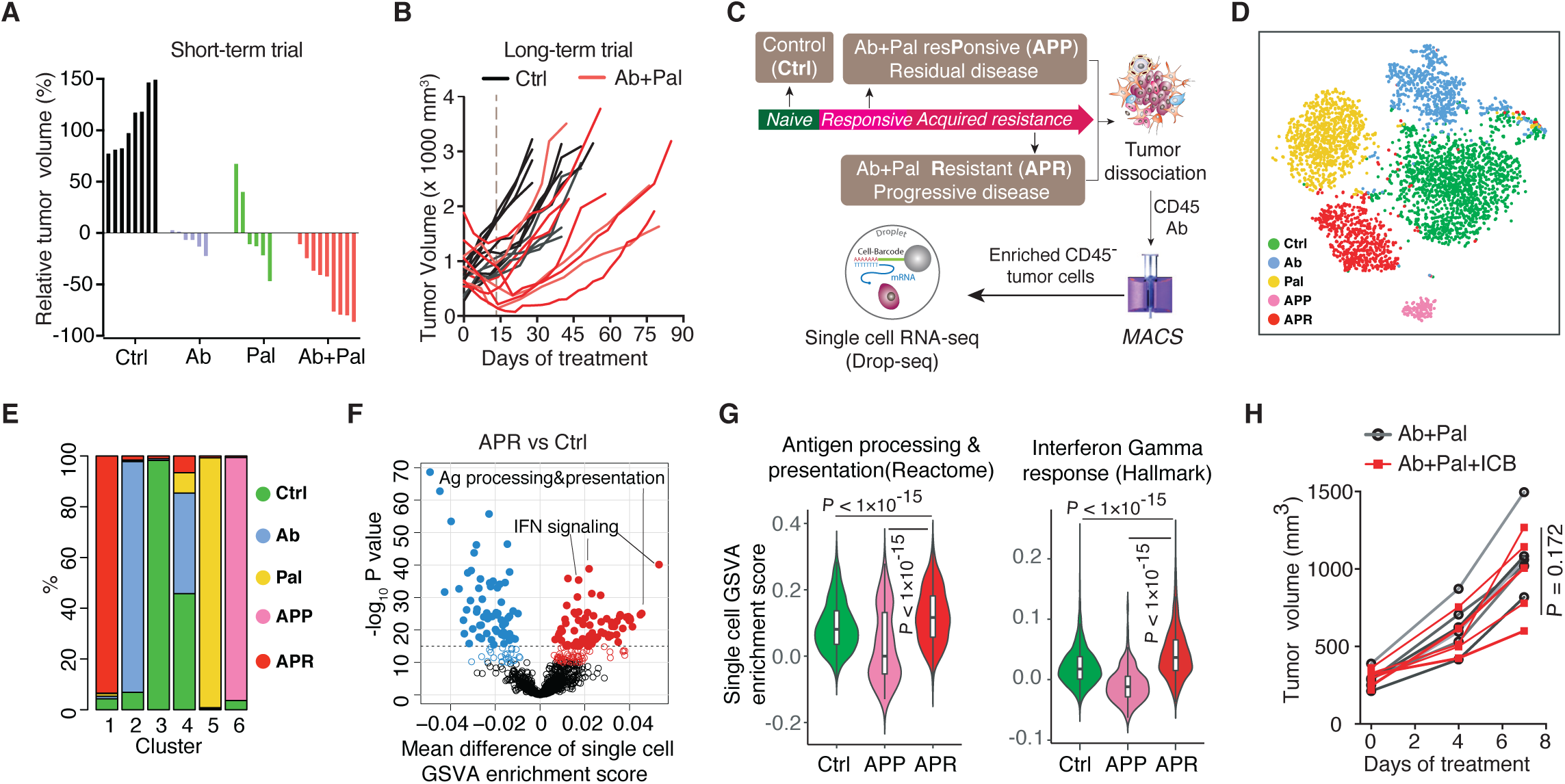
Emergence of resistance to Her2/neu and CDK4/6 combinatorial targeted therapy with increased antigen presentation and IFN signaling in tumor cells. (**A**) Waterfall plots showing percent change of tumor volume with 14-day’s treatment in MMTV-Neu mice (n=8,6,6 and 9 for Ctrl, Ab, Pal and Ab+Pal). Ctrl, vehicle treated; Ab, anti-Her2/Neu antibody; Pal, CDK4/6 inhibitor Palbociclib. (**B**) Tumor volume curves showing tumors rebounded during sustained Ab+Pal combination treatment (n=12 for Ctrl and n=10 for Ab+Pal). (**C**) Schematic for sample processing, enrichment of tumor cells and Drop-seq based single-cell RNA sequencing. (**D**) t-distributed stochastic neighbor embedding (t-SNE) plots colored by treatment groups and clustering of 4711 tumor cells derived from Ctrl, Ab or Pal alone, responsive/residual tumors (APP) and resistant tumors (APR) with Ab+Pal treatment. Each point represents a single cell. (**E**) Abundance of each cell cluster in tumors with indicated treatment as presented and classified in panel D. (**F**) Volcano plots comparing ssGSEA enrichment score of 1053 canonical pathways/gene sets of the C2 collection of Molecular Signatures Database between APR and Ctrl single cell RNA-Seq data. Each point represents one pathway/gene set. X-axis, mean difference of single cell ssGSEA enrichment score; Y-axis, −log10 (P-value by t-test). (**G**) ssGSEA enrichment score violin plots for single cells in each group for indicated signatures. (**H**) Volumes of Ab+Pal resistant tumors after treatment combining immune checkpoint blockades (n=7 for Ab+Pal and n=6 for Ab+Pal+ICB). ICB, anti-CTLA4 and anti-PD-1 antibody cocktail. P-value by two-tailed Student’s t-test.

To explore the molecular underpinnings of the development of resistant to Ab+Pal treatment, we performed single-cell RNA sequencing (scRNA-seq) on enriched tumor cells from control (naive to treatment), treatment responsive (residual disease, 10-14 days with Ab+Pal treatment, APP) and treatment resistant tumors (progressive disease, 45-75 days with Ab+Pal treatment, APR) (Fig. 1C). First, we used nonlinear dimensionality reduction (t-distributed stochastic neighbor embedding, t-SNE) analysis to examine global transcriptional features across tumor cells derived from control, single treatment or Ab+Pal combination treatments (Fig. 1D). We observed distinct distribution patterns of tumor cells among the indicated different treatment tumors, with a total of 6 clusters (Supplemental Fig. 2, A and B). Generally, individual cells derived from different treatments tended to cluster together as distinct clusters (cluster 1, 2, 3, 5, 6), suggesting the transcriptional profiles were greatly influenced by treatments (Fig. 1D and Supplemental Fig. 2, A to C). Cluster 3, 2, 5 and 1 were largely representing cells derived from control, Ab only, Pal only and APR tumors, respectively (Fig. 1, D and E). One exception to the seemingly mutually exclusive clustering based on treatment was cluster 4, which was characterized by the high expression of proliferation genes such as *Top2a, Cdk1, Mki67* and *Cenpa* (Supplemental Fig. 2D). Cells in the cluster 4 were derived from either control, Ab or Pal single treated and combination treated resistant tumors, suggesting that subpopulation of tumor cells conferred tolerance to treatment or adapted to drug selection. Cluster 6 mostly represented the cells from the responsive tumors (residual disease) (Fig. 1, D and E) with high expression of *Ltf, Scd1, Lipa, Lrg1* and *Ifrd1* genes (Supplemental Fig. 2D). Of particular interest is cluster 1, which is predominantly composed of cells from APR tumors (Fig. 1E, Supplemental Fig. 2, B and C) with high expression levels of *Psmb8, Ptpn2, Ifitm3, Map2k3, Iigp1, Irgm1, Cxcl1 and Cxcl2* (Supplemental Fig. 2D). Besides the dominant clustering as cluster 1, APR tumor cells also spread into other clusters, indicating the nature of heterogeneity.

To examine the functional implications of gene signatures unique to each cluster, we performed single-sample gene set enrichment analysis (ssGSEA) focusing on scRNA-seq data derived from control, combination treatment responsive and resistant tumors. We applied canonical pathways from KEGG, REACTOME and BIOCARTA gene sets of the C2 collection of Molecular Signatures Database (MSigDB) to each single cell to obtain enrichment score for each signature, and did pair-wise comparison by *t*-test (Fig. 1F and Supplemental Fig. 2E). Targeting G1-to-S-phase cell-cycle transition of tumor cells is recognized to be the primary mechanism of action of CDK4/6 inhibitors. Gene sets enrichment analysis revealed that, overall, G-S-phase cell-cycle transition and mitotic activity were downregulated in short-term combination treatment responsive (residual) tumors (APP) compared to control treated tumors, while resistant tumors (APR) showed a reprogramed cell-cycle machinery with slight enhanced mitotic activity (Supplemental Fig. 2F), which was consistent with Ki67 staining result (Supplemental Fig. 1, A and E). In the responsive residual tumors (APP), an enrichment of genes involved in both death receptor ‘P75 NTR signaling’ and ‘NFκB is activated and signals survival’ (Supplemental Fig. 2, E and G), suggesting that Ab+Pal treatment induced death signaling and reprogrammed survival signaling to adapt to the treatment. In the resistant tumors (APR), notably, enrichment analysis showed that ‘antigen processing and presentation’ and ‘interferon signaling signatures’ were among the most strikingly differential enriched signatures in the APR tumors compared with control and APP tumors (Fig. 1, F and G, Supplemental Fig. 2 E). Specifically, genes encoding mouse major histocompatibility complex (MHC) class I molecules (*B2m* and *H2d1*), peptide transporters (*Tap1*), proteasome family members for protein degradation and peptide production (e.g. *Psma7, Pamb1, Psmc3* and *Psmd8*) and transporter-MHC interactions (*Tapbp*) exhibited either higher expression levels or more expressing cells in APR tumors (Supplemental Fig. 2H upper panel). Along with the enhanced antigen processing and presentation pathways, signatures including ‘interferon signaling’, ‘interferon gamma response’ (Fig.1, F and G, Supplemental Fig. 2E) and ‘antiviral by IFN stimulated genes’ were also enriched in resistant tumors (Supplemental Fig. 2G). For example, expression of genes involved in innate immune response to viral infection (*Eif2ak2*, double-stranded RNA-activated protein kinase), regulation of interferon signaling (*Ptpn1*), interferon-responsive transcription factors (*Irf7* and *Stat1*) and interferon stimulated/ inducible genes (*Ifi27, Usp18, Xaf1*) were increased in resistant tumors (Supplemental Fig. 2H lower panel). These results at the single-cell transcriptome level indicated that CDK4/6 inhibitor treatment elicits antigen presentation and stimulate IFN signaling, supporting and extending previous observations ^33^. Given that increased antigen presentation and IFN signaling, which suggested an elevated tumor immunogenicity in Ab+Pal resistant (APR) tumors, we next sought to combine immune checkpoint blockades (ICB, anti-CTLA4 and anti-PD-1 antibodies) to overcome or prevent the resistance to Ab+Pal treatment. However, the addition of ICB to the rebound APR tumors showed only modest effect (Fig. 1H, Ab+Pal+ICB), suggesting neither CTLA4 nor PD-1/L1 axis was the major mediator for the resistance. There were likely other factors contributing to the resistant phenotype.

### Single-cell RNA-sequencing reveals distinct immune milieu among different phenotypes and immature myeloid cells are enriched in resistant tumors

We next investigated the TME factors that could potentially mediate the development of resistance. The observation that more CD45^+^ leukocytes in both APP and APR tumors compared to Ctrl (Supplemental Fig. 3) led us to focus on the immune compartment. CD45^+^ tumor infiltrated leukocytes (TILs) were isolated by magnetic-activated cell sorting (MACS) from single-cell suspensions of tumors and single-cell RNA sequencing was performed (Fig. 2A). We obtained a total of 1444 high-quality individual TIL for further analysis (the median number of genes and UMI counts detected per cell was 1523 and 3305, respectively). tSNE clustering identified 9 clusters among these TILs (Fig. 2B, left), based on significant principal components of 3,100 variable genes. Unlike the distribution pattern of tumor cells which were largely dependent on treatment, a great number of TILs from different groups were mixed together or clustered closely (Supplemental Fig. 4A), suggesting their similar transcriptomic properties. Initial examination of top cluster-specific genes revealed major features of macrophage (e.g. *Apoe, Lyz2, C1qb, C1qc*) in cluster 1 (281 cells) and cluster 2 (147 cells) cells (Supplemental Fig. 4B). Cluster 8 (222 cells) and Cluster 9 (92 cells) showed high expression of NK and/or T-cell genes (e.g. *Nkg7, Gramb, Cd3g, Cd3d, Trbc2, Cd8b1*) (Supplemental Fig. 4B). The classification of macrophage, T and NK cells (Fig. 2B, right) was also supported by visualization expression of key marker genes across the single-cell data (Supplemental Fig. 4C). Cluster 3 [50 cells, mostly (38/50) derived from APP] was characterized by *Sparc, Fstl1, Igfbp7, Timp3* and collagen genes, including *Col1a1, Col1a2 and Col3a1*(Supplemental Fig. 4B). Cluster 7 (17 cells all from APP) comprised genes characteristic of both dendritic cell and macrophage, such as *Cd72, Itgal* and *Fcrl5* (Supplemental Fig. 4B). Of note, cells of clusters 4 (327 cells) and 5 (191 cells) displayed high expression of monocyte genes (*Cd14* and *Lcn2*) with the unique expression of *Arg1* and *Xbp1* (Supplemental Fig. 4, B to D), which are molecular features associated with myeloid-derived suppressor cells (MDSCs) ^37,38^. Cluster 6 (117 cells) showed intermediate expression of cluster 1 and 2-specific genes, as well as cluster 4,5-related genes, suggesting that these cells might be an intermediate state between macrophage and cells of cluster 4,5. Therefore, cells of cluster 4, 5 and 6 were annotated as immature myeloid cells (IMCs) (Fig. 2B, right). The above single-cell transcriptome-based profiling and classification of TILs indicated a distinct shift of the immune microenvironment among control, Ab+Pal responsive and resistant tumors. The responsive tumors contained a higher frequency of T and NK cells while immune microenvironment of resistant tumors was dominated by IMCs (Fig. 2C).

**Fig. 2.**
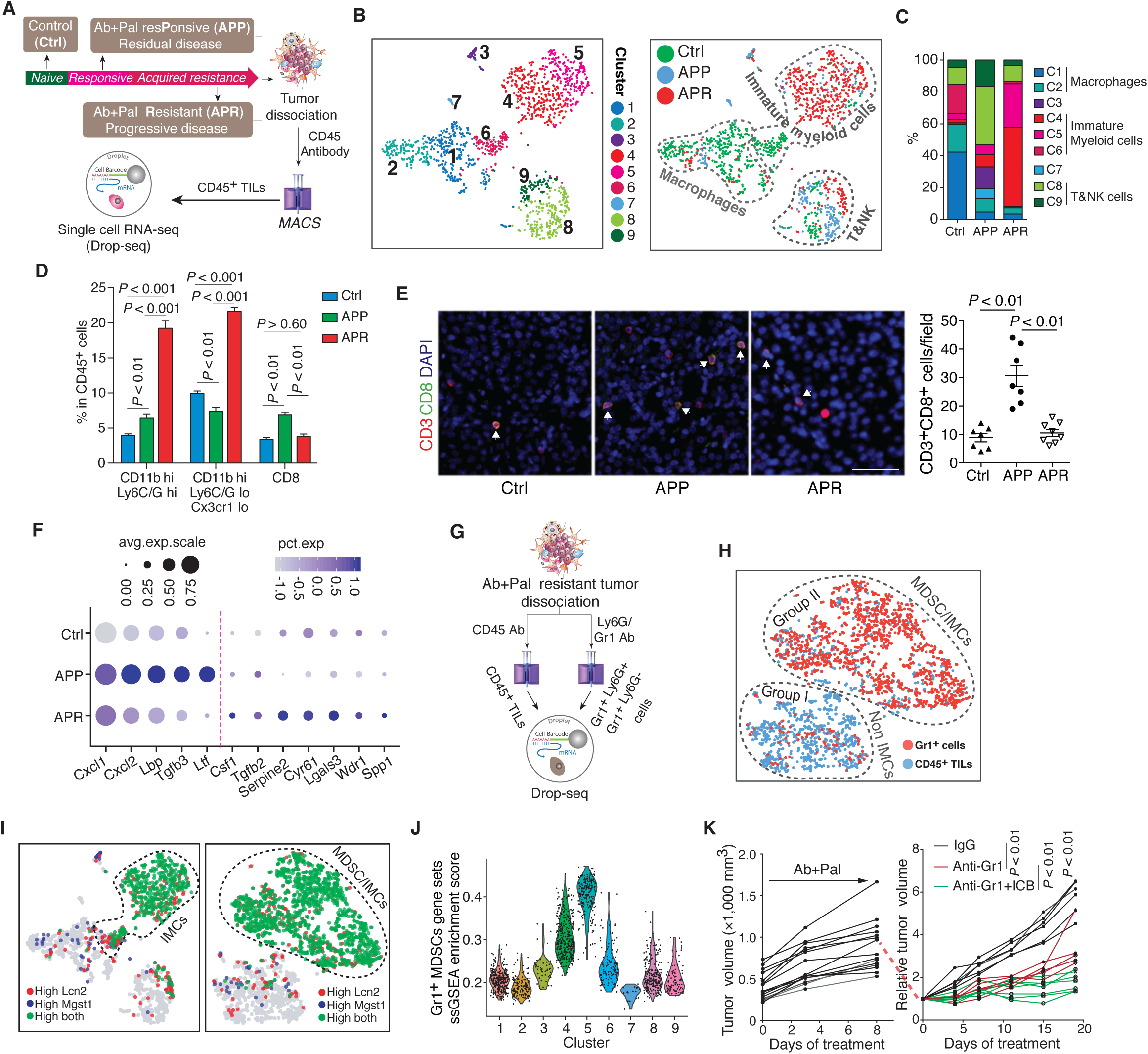
Distinct immune milieu among different phenotypes and immature myeloid cells are enriched in Her2/neu antibody plus palbociclib treatment resistant tumors. (**A**) Schematic for tumor dissociation, isolation of tumor infiltrated leukocytes (TILs) and Drop-seq based single cell RNA sequencing. Ctrl, vehicle treated control; Ab, anti-Her2/Neu antibody; Pal, CDK4/6 inhibitor Palbociclib; TILs, tumor infiltrated leukocytes. (**B**) Clustering of 1444 TILs derived from Ctrl, APP and APR tumors and t-SNE plot colored by clusters (left) and annotation of TIL-clusters on t-SNE plot colored by treatment groups (right). Each point represents a single cell. (**C**) Abundance of each cluster [as clustered and annotated in (B)] in TILs with indicated treatment. (**D**) Analysis of immune cell populations among CD45+ TILs by mass cytometry. Immune cell populations were identified by overlaying the expression of markers on viSNE plots as shown in Fig. S5. (**E**) Representative images and quantification of CD3 and CD8 immunofluorescence staining for Ctrl, APP and APR tumors. Arrows indicate CD3 and CD8 double positive cells. Scale bar, 50 μm. (**F**) Single cell transcriptional analysis of tumor-produced cytokines and chemokines. The size of each circle reflects the percentage of cells in a treatment group where the gene is detected, and the color intensity reflects the average expression level within each treatment group.(**G**) Schematic for tumor dissociation, isolation of CD45+ TILs and Gr1+ cells and Drop-seq based single cell RNA sequencing. (**H**) t-SNE plot of single cell RNA sequencing data from CD45+ TILs and Gr1+ cells. Each point represents a single cell.(**I**) Overlay of marker genes for Gr1+ cell population identified from experiment (F) on t-SNE plots derived from experiment (A) and (F), respectively. (**J**) Violin plot showing distribution of ssGSEA enrichment score among TIL-clusters identified from experiment (A). Experiment (F) generated MDSCs signature (top 300 differential expressed genes for MDSC population) was used for enrichment analysis. Each point represents a single cell. (**K**) Growth of Ab+Pal resistant tumors by adding Gr1 antibody and ICB. Ab+Pal resistant tumors were transplanted to recipient MMTV-Neu mice and first treated with Ab+Pal to acquire the resistance phenotype (left) and then adding IgG (n=6), Anti-Gr1 (n=5) or Anti-Gr1+ICB (n=7) treatments were followed (right). *P-*value in D,E and K by one-way ANOVA with Tukey’s test.

To connect the canonical cell surface markers with the observed transcriptome heterogeneity of TILs, we profiled the TILs of control, APP and APR tumors using CyTOF. CD45^+^ live cells were analyzed using viSNE algorithm (Supplemental Fig. 5, A and B), a data visualization tool that creates a two-dimensional view of high-parameter biological information while conserving the richness of the data at the single-cell level ^39^. We first observed an increase of CD11b^high^ myeloid cells while a decrease of CD11b ^low^ cells in APR tumors compared to those of the control and APP tumors (Supplemental Fig. 5C). Consistent with the trend of scRNA-seq profiling and classification, responsive tumors exhibited more T and NK cells among the infiltrated CD45^+^ cells (Supplemental Fig. 5C). Lymphocyte antigen 6 complex (Ly6C/G) and chemokine (C-X3-C motif) receptor 1 (Cx3cr1) are valuable markers with both phenotypic and functional significance for myeloid cells. Closer examination of CD11b^high^ myeloid cells showed an increase of CD11b^high^ Ly6C/G^high^ (19.23% in APR tumors compared to 3.91% and 6.41% in control and responsive tumors, respectively) and CD11b^high^ Ly6C/G^low^Cx3cr1^low^ (21.63% in resistant tumors compared to 9.94% and 7.41% in control and responsive tumors, respectively) subpopulations in resistant tumors (Fig. 2D and Supplemental Fig. 5D). Of note, CD11b and Ly6C/G (Gr-1) are recognized as phenotypic markers of mouse myeloid-derived suppressor cells (MDSCs). Immunofluorescence staining confirmed a significant decrease of CD8^+^ T-cell infiltration (Fig. 2E) and a great increase of MDSCs (Supplemental Fig. 5F) in the resistant tumors compared to responsive tumors. Collectively, these observations revealed that APP tumors were infiltrated with more T and NK cells while, in contrast, APR TME were dominated by IMCs.

The dominant presence of IMCs suggested an immunosuppressive microenvironment in the resistant tumors. To understand the possible mechanisms involved in the transition of the immune microenvironment, the effect of Ab+Pal treatment on expression of cytokines and chemokines was investigated, as tumor-produced factors are critical for the recruitment and functional properties of TILs ^14^. Single cell transcriptional analysis of tumor cells revealed that several secreted factors involved in recruitment or chemotaxis of myeloid cells were increased, including *Cxcl1, Cxcl2, Tgf*β*3* and lactotransferrin (*Ltf*) after short-term Ab+Pal treatment (Fig. 2F). As an immunoregulatory factor, Ltf has been reported as a driver for accumulation and acquisition of immunosuppressive activity of MDSCs ^40^. On the other hand, expression of multiple cytokines and chemokines associated with myeloid cell recruitment and differentiation, including *Csf1, Tgf*β*2, Serpine2, Cyr61* and *Lgals3* were upregulated in Ab+Pal resistant tumor cells (Fig. 2F). It has been shown that colony stimulating factor 1 (CSF-1) is important for development and activation of MDSCs ^37,38^. These data indicate that tumor cells are capable of evolution/adaptation through the production of multiple immunomodulatory factors to establish an immunosuppressive environment to acquire and sustain resistance to Ab+Pal combination treatment.

### Single-cell transcriptome profiling annotated immature myeloid cells shared molecular characteristics of myeloid-derived suppressor cells

Noticeable in APR tumors, the infiltrating immature myeloid cells (IMCs) characterized by scRNA-seq (clusters 4 and 5) possessed certain molecular characteristics of MDSCs. This observation led us to explore the potential association between the transcriptome profiling identified IMCs and the surface markers defined MDSCs through transcriptomic analysis. We employed a tumor transplantation model by transplanting APR tumors to mammary fat pads of recipient syngeneic MMTV-Neu mice (12 to14-week old). After forming palpable tumors, the recipient mice were treated with Ab+Pal for 1-3 weeks to establish/ensure the resistance phenotype. First, we isolated tumor infiltrated Gr-1+ cells (including Gr1^high^Ly6G^+^ and Gr1^dim^Ly6G^-^ populations) and found that these cells inhibited the proliferation of CD4+ and CD8+ T cells *in vitro* (Supplemental Fig. 6A), an important functional characteristic of MDSCs ^37^. Next, Gr-1+ cells and CD45+ TILs were isolated in parallel from the transplanted Ab+Pal resistant tumors and scRNA-seq was performed (Fig. 2G). 2,471 cells in total (with 1,318 Gr-1+ cells and 1,153 CD45+ TILs) were analyzed after quality control filtering. Unsupervised clustering separated these heterogeneous cells into two apparent subgroups: one group was predominantly from CD45+ TILs (group I) while the other group of cells (group II) were dominated by Gr1^high^Ly6G^+^ and Gr1^dim^Ly6G^-^ cells (Supplemental Fig. 6B). Group I cells showed high expression of macrophage genes (*CD14, Emr1* or *F4/80, Apoe, Lyz2*) and T/NK cell-related genes (*Cd3e, Nkg7, Cd4* and *Cd8a*), while group II cells exhibited enriched expression of MDSC related genes, *Arg1* and *Xbp1* (Supplemental Fig. 6C). Thus, these two groups of cells were annotated as non-IMCs and MDSC/IMCs, respectively (Fig. 2H). Based on marker genes of group II cells, we generated Gr-1+ MDSCs signature (Supplementary Table 1). We found that *Lcn2* and *Mgst1*, two of the marker genes of Gr-1+ MDSCs identified by scRNA-seq analysis, were also specifically present in previously identified IMCs related cells (Fig. 2I). Indeed, flow sorting and qPCR of APR tumors showed significant higher expression levels of both *Lcn2* and *Mgst1* in Gr1^+^ cells compared to T cells and macrophages (Supplemental Fig. 6D). Further, single-cell gene set enrichment analysis using our custom experimentally generated Gr-1^+^ MDSCs signature revealed that the geneset was also enriched in the annotated IMCs, particularly in cluster 5 cells (Fig. 2J). This analysis demonstrated that transcriptomic profiling identified IMCs (predominately presented in the APR tumors) displayed similar transcriptome profiles to previously defined Gr-1+ MDSCs. Defined by cell surface marker expressions, the MDSCs have been sub-grouped as Gr1^high^Ly6G^+^ and Gr1^dim^Ly6G^-^ MDSC, which largely reflect granulocytic/polymorphonuclear and monocytic lineage of MDSCs ^37,38^. Interestingly, based on the single cell transcriptome profiles, in our case, the Gr1^high^Ly6G^+^ and Gr1^dim^Ly6G^-^ cells were clustered closely or mixed together (Supplemental Fig. 6B), suggesting the transcriptional similarity between these two groups, despite the distinct cell surface marker differences.

### Depletion of IMCs sensitizes Ab+Pal resistant tumors to ICB treatment

We next assessed whether the increased Gr-1+ MDSCs population was functionally important for Ab+Pal resistant phenotype. After confirming the resistance phenotype of transplanted APR tumors (Fig. 2K, left), the mice were further treated with either anti-Gr1 antibody or anti-Gr1 plus ICB. MDSCs depletion with anti-Gr1 antibody inhibited growth of APR tumors (Fig. 2K, right), suggesting a pro-tumor role of Gr1+ MDSC cells. Notably, addition of ICB showed further enhanced tumor inhibitory effect (Fig. 2K, right), indicating that MDSCs were not only involved in promoting APR phenotype but also in hindering maximal efficacy of ICB.

### Identification and selection of cabozantinib as a potential IMCs targeting drug

Motivated by the above results, we sought to modulate or target IMCs in APR tumor to overcome Ab+Pal resistance. With a goal of potentially repurposing existing drugs to combat the resistance, we screened the drug target portfolios of FDA-approved small molecular protein kinase inhibitors (PKIs) against the single-cell transcriptome of TILs. We observed that in addition to EGFR and/or HER2 inhibitors, cabozantinib target genes (including *Met, Kit, Axl, Kdr/Vegfr2, Flt3*) and Lenvatinib target genes (including *Vegfr1/2/3, Pdgfr, Fgfr, Kit, Ret*) were significantly enriched in TILs from APR tumors compared with those of responsive and control tumors (Supplemental Fig. 7, A and B). Cabozantinib (Cabo), an orally bioavailable tyrosine kinase inhibitor, is approved for metastatic medullary thyroid cancer and renal cell carcinoma. Cabo also showed promising clinical activity for metastatic breast cancer in a phase 2 trial ^41^ and is being further investigated (ClinicalTrials.gov NCT01441947 and NCT02260531). This prompted us to conduct an in-depth examination of Cabo. Unlike TILs (Fig.3A, left panel), the enrichment of Cabo target genes in APR tumor cells showed no significant changes compared to control and APP tumor cells (Supplemental Fig. 7C). Interestingly, IMCs showed the highest average enrichment score (Fig. 3A right panel) and clusters 4, 5 (IMC clusters) possessed more cells with a relatively high enrichment score (Supplemental Fig. 7D). Specifically, IMC clusters contained a higher percentage of Kit and/or Met expressing cells compared to either T&NK cells or macrophages (Fig. 3B). Moreover, the IMC population derived from APR tumors were largely composed of Kit and/or Met expressing IMCs (Fig. 3C). Consistently, Gr1+ MDSC/IMCs isolated from APR tumors also showed much higher percentage of Kit and/or Met expressing cells than other non-IMCs (including macrophages, NK and T cells) (Supplemental Fig. 7, E and F). Indeed, qPCR confirmed higher expression levels of *Kit* and *Met* in CD45^+^ TILs from APR tumors compared to those of from either Ctrl or APP tumors (Supplemental Fig. 7G). In addition, Gr1+ MDSC/IMC population showed the highest expression of *Kit* and *Met* among the sorted cell types (Supplemental Fig. 7H). Altogether, our single-cell transcriptome profiling analysis and qPCR validation suggested that IMCs in APR tumors might be targetable by Cabo.

**Fig. 3.**
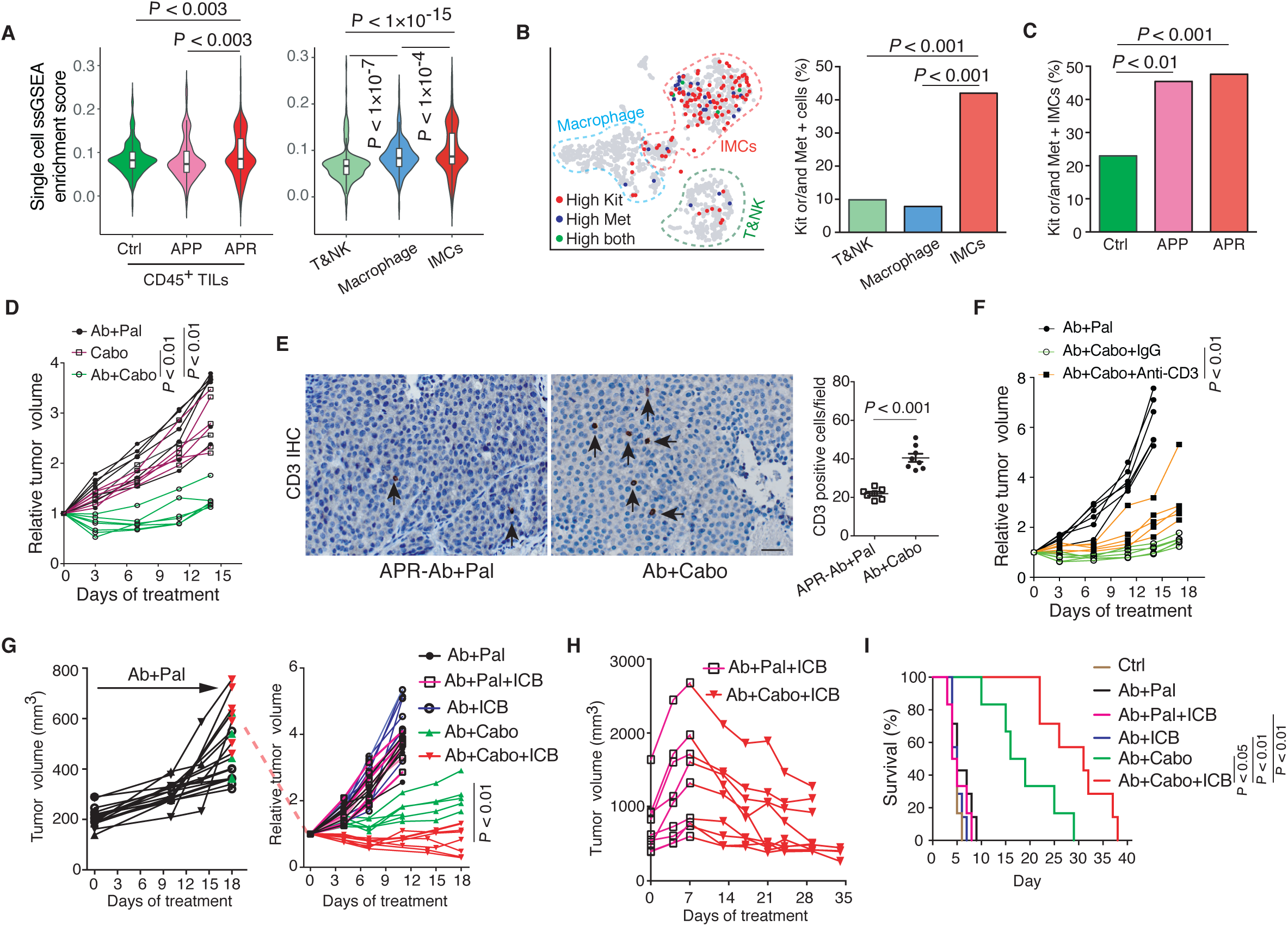
Identification and evaluation of single-cell RNA-seq analysis-driven therapeutic strategy for resistant tumors to Ab+Pal treatment. **(A)** Enrichment analysis of cabozantinib target genes across single TILs grouped and plotted by different phenotypes (left) or by different immune cell types (right) as annotated in Fig.2 (B). *P*-value by Student’s t-test (two-tailed). **(B)** Expression distribution of Kit and Met on t-SNE plot (left) and quantification of Kit and/or Met expressing cells among different immune celltypes as annotated (right). *P*-value by three-sample Chi-square test. **(C)** Abundance of Kit and/or Met expressing IMCs among tumors with different phenotypes. *P*-value by three-sample Chi-square test. **(D)** Growth of Ab+Pal resistant tumors with Ab+Cabo treatment (n=7 for Ab+Pal, n=6 for Cabo and n=7 for Ab+Cabo). **(E)** Representative images and quantification of CD3 immunohistochemistry staining for tumors with Ab+Pal or Ab+Cabo treatment. Scale bar, 20 μm. *P*-value by Student’s t-test. **(F)** T-cell depletion during Ab+Cabo treatment against Ab+Pal resistant tumors. *P*-value by Student’s t-test. **(G)** Relative volumes of Ab+Pal resistant tumors after treatment with Cabo and ICB. Ab+Pal resistant tumors were transplanted to recipient MMTV-Neu mice and first treated with Ab+Pal to acquire the resistance phenotype (left), then treated with Ab+Pal (n=6), Ab+Pal+ICB (n=6), Ab+ICB (n=8), Ab+Cabo (n=5), or Ab+Cabo+ICB (n=7) for 2 weeks (right). **(H)** Growth of Ab+Pal resistant tumors after sequential treatment with Ab+Pal+ICB and Ab+Cabo+ICB (n=9). Ab+Pal resistant tumors were first treated with Ab+Pal+ICB for 1 week then switched to Ab+Cabo+ICB treatment for 3 weeks. **(I)** Survival time to doubled tumor volume from experiment in (D). *P*-value by log-rank (Mantel-Cox) test. Cabo, protein kinase inhibitor cabozantinib. ICB, immune checkpoint blockades cocktail with anti-CTLA4 and anti-PD-1 mAb.

### Evaluation of single-cell RNA-seq analysis-driven therapeutic strategy for Ab+Pal resistance

To evaluate the effectiveness of Cabo, a potential MDSC/IMCs targeting inhibitor, for treating APR tumors, we again employed the transplantation model similar to previous experiments (Fig. 2K) to establish a cohort of mice with relatively uniform tumors. The transplanted APR tumor bearing mice were either continuously treated with Ab+Pal or with Ab+Cabo. Although Cabo monotherapy at the given dose had no anti-tumor activity, Ab+Cabo treatment significantly inhibited tumor growth (Fig. 3D). Interestingly, Ab+Cabo treated tumors showed increased T cell infiltration compared to tumors with continuous Ab+Pal treatment (Fig. 3E) and T cell depletion during Ab+Cabo treatment resulted in significant reduction of tumor suppression (Fig. 3F), suggesting that the optimal therapeutic activity of Ab+Cabo against APR tumors is dependent on T cells. Next, addition of ICB to Ab+Cabo combination further improved therapeutic efficacy (Fig. 3G). Consistent with our previous observation as shown in Fig. 1H, Ab+Pal+ICB had limited efficacy on APR tumors (Fig. 3G). Histology analysis of treated tumors revealed a significant increase of tissue hypocellularity (Supplemental Fig. 8A) and a reduced tumor proliferation (Supplemental Fig. 8B) in Ab+Cabo and Ab+Cabo+ICB treated group. Furthermore, in another cohort of mice, although the addition of ICB (Ab+Pal+ICB) had limited effect on APR tumor progression, notably, switching to Ab+Cabo+ICB combination treatment not only blocked tumor progression, but also led to tumor shrinkage (Fig. 3H and Supplemental Fig. 8C). Importantly, both Ab+Cabo and Ab+Cabo+ICB treatment greatly extended survival (time to doubled tumor volume) from a median of ∼5 days in Ctrl and continuous Ab+Pal treatment group to 17.5 days in Ab+Cabo treated group and up to 31 days in Ab+Cabo+ICB treated group (Fig. 3I). Together, these data indicated that Ab+Cabo combination, identified by single-cell transcriptome analysis, was effective in overcoming Ab+Pal resistance, and the application of immunotherapy using ICB further enhanced the anti-tumor activity.

### Cabo and ICB combination subverted immunosuppressive tumor microenvironment and enhanced anti-tumor immune response

It has been previously shown that cabozantinib could synergize with immune checkpoint blockade by attenuating MDSC frequency and immunosuppressive activity in a mouse model of metastatic castration-resistant prostate cancer ^42^. Since Cabo alone did not effectively suppressed the APR tumor growth (Fig. 3D), we speculated that the anti-tumor effect of Cabo-containing combinatorial regimen might be due in part to its activity on modulating IMC in the TME. Thus, the impact of Cabo-containing combination on the immune microenvironment was examined. First, we performed CyTOF analysis focusing on CD45^+^ TILs from APR transplants with either continuous Ab+Pal treatment, Ab+Cabo or Ab+Cabo+ICB combination. Both CD11b^high^ Ly6C/G^high^ and CD11b^high^ Ly6C/G^low^Cx3cr1^low^ populations, which were enriched most significantly in the APR tumors as shown in Fig. 2D, were greatly decreased after Ab+Cabo combination treatment (Fig. 4A and Supplemental Fig. 9). The addition of ICB led to further reduction of the CD11b^high^ Ly6C/G^low^Cx3cr1^low^ population (Fig. 4A). Meanwhile, Ab+Cabo combination treatment showed a mild or moderate increase of CD11b^low^ populations and NK, CD4^+^ and CD8^+^ T cells (Fig. 4A and Supplemental Fig. 9).

**Fig. 4.**
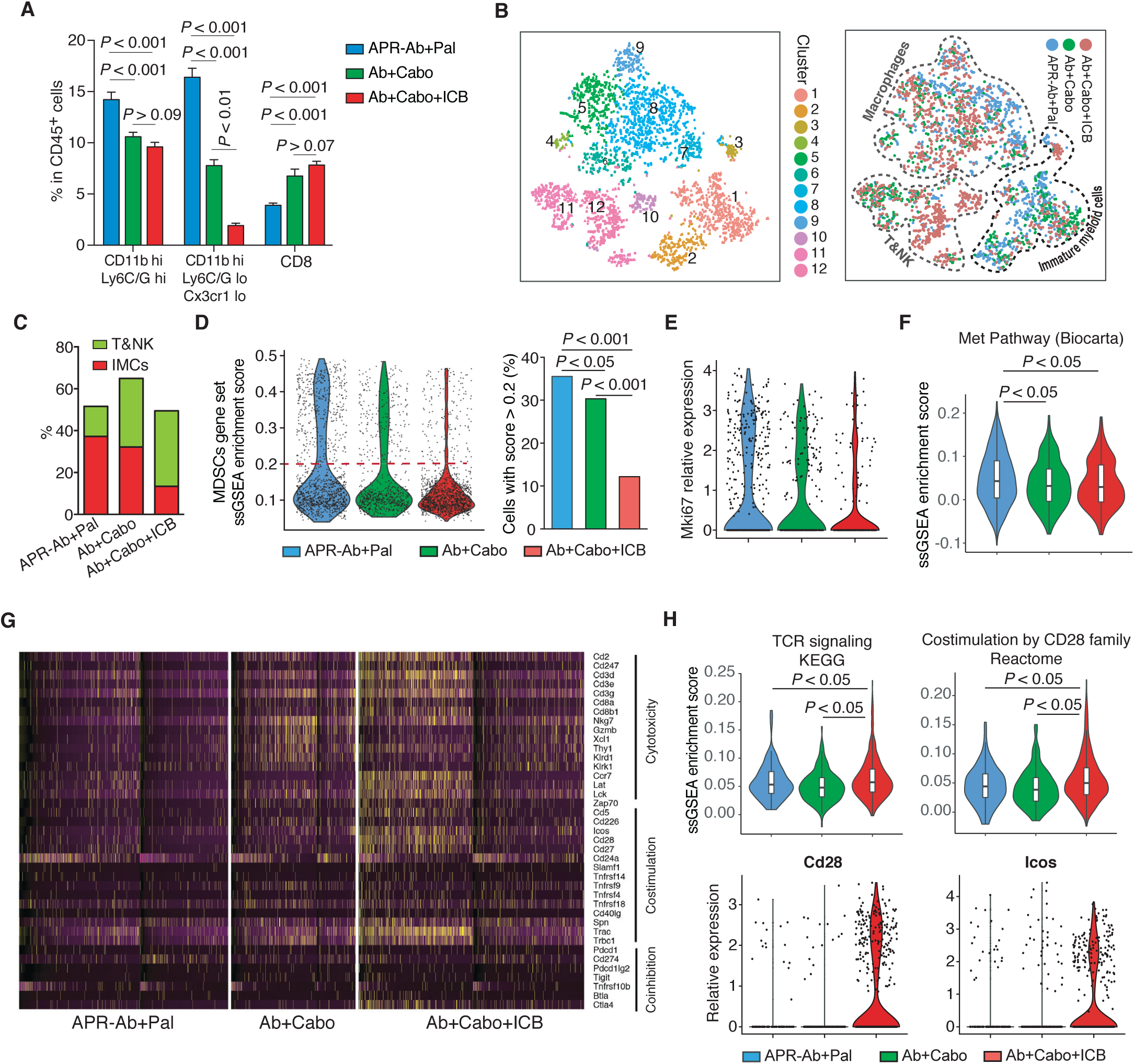
Cabo and ICB combination subverted immunosuppressive tumor microenvironment and enhanced anti-tumor immune response. (**A**) CyTOF characterization of tumor infiltrated immune cell populations and their relative abundance among CD45+ TILs after treatment with Cabo and ICB. *P*-value by one-way ANOVA with Tukey’s test. (**B**) Clustering and annotation of single cell RNA sequencing data including 3168 TILs derived from resistant tumors with continuous Ab+Pal, Ab+Cabo or Ab+Cabo+ICB treatment. left, t-SNE plot colored by clusters; right, annotation of TIL-clusters on t-SNE plot colored by treatment groups. Each point represents a single cell. (**C**) Abundance of T and NK cell and IMC clusters [as clustered and annotated in (B)] in the TILs with indicated treatment. (**D**) Distribution of MDSCs related signature enrichment score (left) and proportion of cells with high enrichment score in the TILs with indicated treatment (right). MDSCs related signature was generated from experiment Fig.2F (top 300 differential expressed genes of MDSC population) was used for enrichment analysis. Each point represents a single cell. *P*-value by three-sample Chi-square test. (**E**) Expression of Ki67 in IMC population [as clustered and annotated in (B)] from tumors with indicated treatment. (**F**) Enrichment of ‘Met pathway’ in IMC population [as clustered and annotated in (B)] from tumors with indicated treatment. (**G**) Heatmap of T cell response signature genes across CD45+TILs. Specific genes from gene sets for T-cell cytotoxicity, costimulation and coinhibition were shown. Each column represents a single cell. (**H**) Enrichment scores for ‘T-cell receptor signaling’ and ‘Costimulation by the CD28 family’ (upper panel) and expression of CD28 and ICOS (lower panel) in T&NK cell population as clustered and annotated in (B). Each point represents a single cell. *P*-value in F and H by two tailed Student’s t-test.

To gain deeper insight into how Cabo and ICB combination modulated the immune response, we next performed scRNA-seq on CD45^+^ TILs. Unsupervised graph-based clustering identified 12 clusters among 3168 TILs from these 3 different treatment groups (including 1185 TILs from tumors with continuous Ab+Pal treatment, 705 Ab+Cabo TILs and 1278 Ab+Cabo+ICB TILs) (Fig. 4B, left and Supplemental Fig. 10A). CD45^+^ TILs derived from different treatment groups were clustered together primarily based on cellular identity rather than treatment (Supplemental Fig. 10A). Examining the expression of immune lineage marker genes, including *Cd14, Csf1r, Cd3e, Nkg7, Cd8a, Cd4 and Lcn2* (Supplemental Fig. 10B) identified three major categories: IMCs (clusters 1-3), macrophages (clusters 4-9) and T and NK cells (clusters 10-12) (Fig. 4B, right). In line with the CyTOF results, this classification of TILs indicated that Ab+Cabo combination treatment decreased tumor infiltrated IMCs and increased T and NK cells (Fig. 4C). Addition of ICB led to further reduction of IMCs population concomitant with mild increase of T and NK cells (Fig. 4C) and IF staining confirmed further increase of CD8^+^ T-cell (Supplemental Fig. 10C). Meanwhile, single cell gene set enrichment analysis showed that our experimentally generated Gr-1+ MDSCs-related signature was decreased among TILs derived from APR-tumors treated with Cabo-containing regimen (Fig. 4D). Particularly, TILs from Ab+Cabo+ICB triple combination exhibited lowest proportion of cells expressing MDSCs signature (Fig. 4D, bar graph). Notably, Cabo-containing regimen suppressed proliferation of IMCs as indicated by Ki67 expression in IMCs (Fig. 4E). Moreover, combination treatment with Cabo attenuated the enrichment of its target genes among IMCs (Supplemental Fig. 10D) and decreased Kit or/and Met expression IMCs (Supplemental Fig. 10E) and Met signaling as well (Fig. 4F). In addition, compared to continuous Ab+Pal treatment, Ab+Cabo or Ab+Cabo+ICB treatment not only promoted infiltration of T and NK cells into the tumors (Fig. 4, A to C and Supplemental Fig. 10C), but also enhanced T-cell related anti-tumor activity, and to a greater extent within tumors after Ab+Cabo+ICB treatment (Fig. 4G, genes associated with cytotoxicity and costimulation were upregulated in Ab+Cabo+ICB-treated TILs). Specifically, pairwise comparison revealed that the enrichment score of “T-cell receptor signaling” and “Costimulation by the CD28 family” signatures across T&NK cell clusters with Ab+Cabo+ICB treatment were higher than those of with Ab+Cabo treatment (Fig. 4H), indicating enhanced T-cell response by ICB, which subsequently increased therapeutic effect. IF staining showed more Granzyme B^+^ CD8^+^ T-cell in Ab+Cabo treated tumors compared to continuous Ab+Pal treatment and additional ICB treatment (Ab+Cabo+ICB) resulted in further increase of Granzyme B^+^ CD8^+^ T-cell (Supplemental Fig. 10F).

### Sequential combinatorial immunotherapy enabled sustained response and significantly prolonged survival of rapidly evolving HER2/neu-positive breast cancers

Our results have shown that Ab+Pal combination treatment initially inhibited spontaneous late-stage HER2/neu-positive mammary tumor. However, resistance to Ab+Pal combination emerged in a short period (Fig. 1B). We found increased immunogenicity (with enhanced antigen presentation and interferon signaling) in tumor cells along with distinct immunosuppressive immature myeloid cells infiltrated in the Ab+Pal resistant TME (Fig. 2). Ab+Pal resistance could be effectively overcome by switching to Cabo-containing combinatorial immunotherapy, which reduced immature myeloid cells and enhanced anti-tumor immunity. These results prompted us to hypothesize that sequential administration of Ab+Cabo (AbC) or Ab+Cabo+ICB (AbC+ICB) combination after a short period of Ab+Pal (AbP) treatment (anti-tumor immunity priming) before the emergence of resistance might achieve a better therapeutic efficacy and prolonged control of tumor progression. To this end, for the control arms, MMTV-neu mice bearing spontaneous advanced tumor (size > 500mm^3^) were continuously treated with either AbP, AbC or AbC+ICB for four weeks (Fig. 5A). And in the sequential treatment group, the MMTV-neu tumor-bearing mice were first treated with AbP for one week for priming of anti-tumor immunity, then switched to AbC or AbC+ICB treatment for another three weeks (Fig. 5A). We observed that sequential regimen with AbC increased progression free survival (PFS) (median of 43 days, P=0.0038) compared with AbP continuously treated mice (median of 29 days). Continuous triple combination regimen (AbC+ICB) without the priming stage exhibited comparable PFS (median of 44 days) to that of sequential AbP+AbC treatment. Notably, prior treatment of AbP priming followed by a sequential combinatorial immunotherapy regimen (AbP/AbC+ICB) further increased PFS (median of 53 days, P=0.0016 vs sequential AbP/AbC, P= 0.025 vs AbC+ICB) significantly (Fig. 5B). This result suggests that AbP priming is important to recondition the tumor immune microenvironment which makes the tumor more sensitive to AbC+ICB combinatorial immune therapy.

**Fig. 5.**
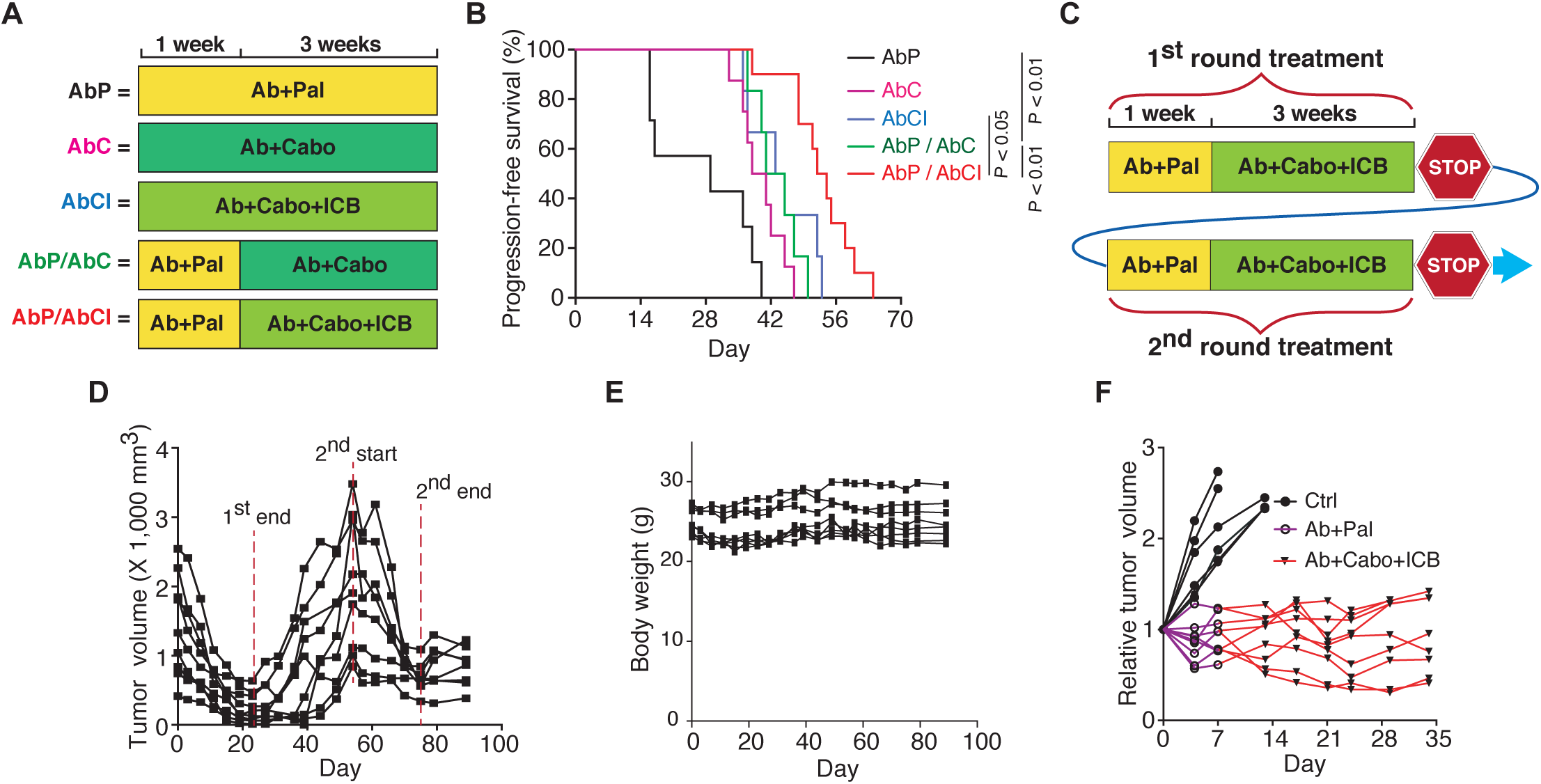
Sequential combinatorial immunotherapy enabled sustained response and significantly prolonged survival of rapidly evolving Her2/neu-positive breast cancer. **(A)**Short-term experimental design for testing efficacy of indicated treatment schedule in MMTV-Neu mice with spontaneous, advanced-stage tumors. (**B**) Kaplan-Meier progression free (without tumor volume increase) survival curves for different treatment and schedule as indicated in (B) (n=6-8). *P*-value by log-rank (Mantel-Cox) test. (**C**) Long-term experimental design for testing efficacy of sequential combinatorial immunotargeted therapy in MMTV-Neu mice with spontaneous, advanced-stage tumors. (**D**) Tumor volume curves of sequential combinatorial immunotargeted therapy (n = 9) as indicated in (C). (**E**) Body weight measurements during treatment as indicated in (D). (**F**) The residual tumors after second round sequential treatment as indicated in (D) were transplanted to recipient syngeneic MMTV-Neu mice (n=6 for Ctrl and n=9 for sequential treatment group). Relative tumor volumes of another round of sequential combinatorial immunotargeted therapy were shown.

Clinically undetectable residual tumors might gradually rebound upon discontinuation of the treatment, which imposes a significant clinical challenge. Encouraged by the significant therapeutic efficacy of AbP/AbC+ICB sequential regimen in inhibiting extremely aggressive APR tumors, we next sought to model the clinical scenario of residual disease and test whether the rebounded-tumors will acquire resistance to the sequential combinatorial immunotherapy (Fig. 5C). Strikingly, a second round of sequential combinatorial immunotherapy was almost as effective as the first round of AbP/AbC+ICB in shrinking the rebounded-tumors (Fig. 5D). Throughout two courses of treatment, sequential combinatorial treatment (AbP/AbC+ICB) was well tolerated and there was no significant weight loss observed (Fig. 5E). To further explore the sustainability of the sequential combinatorial regimen in controlling the tumor relapse, we transplanted the residual tumors after the second-round of sequential treatment to a cohort of recipient syngeneic MMTV-Neu mice. Compared to controls, sequential combinatorial treatment (AbP/AbC+ICB) continuously to inhibit tumor progression during the 3rd round of treatment (Fig. 5F), enabling a sustained control of the extremely aggressive tumors.

## Discussion

Small molecule CDK4/6 inhibitors are one of the most exciting classes of targeted therapies in treating ER-positive breast cancers ^43,44^. Besides, recent studies have demonstrated an interplay between HER2 signaling and cell cycle machinery. Targeting cyclin D1-CDK4/6 complex sensitizes HER2+ tumors to anti-HER2 treatment through cyclin D1 and mTOR pathway cross-talk ^45,46^. These exciting pre-clinical data warranted clinical proposition of CDK4/6 inhibitors to patients with HER2+ breast cancer. Multiple new clinical trials are currently being conducted to examine the clinical efficacy of CDK4/6 inhibitors in advanced HER2+ breast cancer patients ^35,36^. For example, the MonarcHER study (ClinicalTrials.gov identifier: NCT02675231) aims to evaluate efficacy of abemaciclib in treating patients with locally advanced or metastatic HER2+ breast cancer after prior exposure anti-HER2 therapies. Another global phase III PATINA study (ClinicalTrials.gov identifier: NCT02947685) examines the benefits of adding palbociclib in enhancing therapeutic efficacy of current anti-HER2 therapy. Results from the PATRICIA trial (ClinicalTrials.gov Identifier: NCT02448420), a phase II study of palbociclib and trastuzumab with or without anti-ER treatment in HER2+ metastatic breast cancers, suggested that luminal subtype breast cancer correlates with a better progression-free survival compared to non-luminal tumors ^47^. In line with prior pre-clinical studies and recent clinical observations, in our work, we observed similar synergistic benefit of combining CDK4/6 inhibitor palbociclib with anti-HER2 therapy in the classic MMTV-neu mouse model (Fig. 1A). Unfortunately, despite the initial response to palbociclib plus anti-HER2 antibody (Ab+Pal group), the residual tumor under this combinatorial treatment quickly developed resistance and regained the uncontrollable tumor growth (Fig. 1B). While acquisition of resistance to new targeted therapy regimen is not completely unexpected, in light of current clinical trial efforts in using CDK4/6 inhibitors, this rapid development of resistance to palbociclib plus anti-HER2 therapy is concerning. An understanding of the potential mechanisms of drug resistance/inefficacy and the exploration of alternative therapeutic strategies is urgent and critical.

To dissect the cellular and molecular underpinnings of such a fast-evolving drug resistance phenotype, in the present study, we employed single-cell RNA-seq approach to characterize tumors at different stages - from naive untreated stage (Ctrl) to initial response/residual disease stage (APP) and resistant stage (APR). Although one of the most significant transcriptome changes in the APR tumor cells is the up-regulation of IFN signaling and antigen presentation, implying tumor immunogenicity elicited by Ab+Pal treatment (Fig. 1F and G), adding immune checkpoint blockade (ICB) had only a limited effect in inhibiting APR tumors (Fig. 1H). Interestingly, we found that the APR phenotype associated with a significant accumulation of immunosuppressive immature myeloid cell in the TME (Fig. 2), which could be effectively targeted by cabozantinib as identified by single-cell RNA-seq profiling (Fig. 3). We observed that combinatorial immunotherapy by targeting immature myeloid cells and enhancing T-cell activity concomitantly could cooperate to exert optimal therapeutic efficacy in the treatment of aggressive resistant tumors (Fig. 3 and Fig.4). Furthermore, we demonstrated that sequential combinatorial immunotherapy enabled a sustained response and significantly improved outcomes of rapidly evolving HER2/neu-positive breast cancer (Fig. 5).

At the transcriptomal level, the immature myeloid cells identified in our study greatly resemble myeloid-derived suppressor cells (MDSCs). MDSCs represent a heterogeneous population of largely immature myeloid cells with an immune suppressive activity. Two major subsets (monocytic and polymorphonuclear MDSC) have been identified and characterized ^37,38^. However, the current characterization of MDSC relies mostly on the functional level (e.g. ex vivo T-cell suppression assays). The specific and reliable molecular features contributing to the function of MDSCs, especially under the drug resistance context, have not been well defined at the single cell level. Defining the mechanistic underpinnings that drive MDSC phenotypes and their immune suppressive properties in tumors is essential for the development of MDSC-specific therapeutic interventions. In this study, we performed single cell transcriptomic analysis of Gr1^high^Ly6G^+^ and Gr1^dim^Ly6G^-^ cells derived from the APR tumors. Interestingly, the Gr1^high^Ly6G^+^ and Gr1^dim^Ly6G^-^ cells, which are believed to represent granulocytic/polymorphonuclear and monocytic lineage of MDSCs, to a great extent, displayed transcriptomic similarity under the APR context. These similarity may reflect common features of these heterogeneous and plastic myeloid cell subsets in the APR tumor context in suppressing anti-tumor immunity ^48–50^. Furthermore, guided by the single-cell transcriptome signatures, targeting immature myeloid cells by switching to cabozantinib, a clinically actionable strategy, suppressed APR tumors and sensitized tumor to ICB (Fig. 3 and Fig. 5). Since there are a number of on-going clinical studies evaluating HER2 and CDK4/6 co-targeting in breast cancer patients, our study might be valuable in guiding future clinical practice to overcome potential emerging therapeutic resistance. Given the abundance of immature myeloid cells in APR tumors and their apparent tumor-promoting functions, targeting or modulating those cells, which have a relatively stable genome compared to cancer cells, is a clinically appealing strategy. We envision that the signature of MDSCs generated from our study may shed more light on the molecular underpinning of immature myeloid cells and guide the development of therapeutic interventions to precisely target MDSCs.

In this study, we demonstrated that identifying TME changes occurring after treatment is essential for designing more effective combinatorial regimen to combat the tumor evolution. We believe targeting these TME changes early in the disease progression/development of resistance, rather than after outright resistance, will deliver better clinical outcomes, since even short-term drug treatment induces phenotypes promoting drug resistance. Traditionally, targeted therapy is not effective once the tumor develops new mutation or engages alternative pathways to circumvent the drug target on tumor cells. In contrast to the above notion, one of the promises of immunotherapy with ICB is to deliver durable responses in disparate tumor types by reinvigorating antitumor immunity. Our results showed that long-term anti-HER2/neu antibody and CDK4/6 inhibitor combination treatment led to significant increase of immunosuppressive immature myeloid cells in the TME, which in turn diminished the efficacy of ICB. In order to maximize the utility of ICB in treating breast cancer, especially APR resistant tumors, define immune suppressive components and rationally designed combinatorial regimen to provoke tumor immunogenicity is indispensable. In this study, we have demonstrated that sequential administration combinatorial targeted therapy with additional immunotherapy could deliver a durable therapeutic efficacy, thus leading to prolonged stable disease and less drug resistance for the rapidly evolving HER2/neu positive breast cancer. This long-lasting disease control by such treatment might also engage or enhance immune memory, considering the optimal therapeutic activity of Ab+Cabo with a dependence on T cells. These findings provide insights into how and when to optimally integrate immunotherapies against even aggressive breast cancer with extensive prior treatments.

In the present study, although cabozantinib-containing regimen inhibit immature myeloid cells in the APR tumor (Fig. 4), we could not exclude the possible direct effect of cabozantinib on tumor cells and other stroma compartments. Our findings provided clues as to how immature myeloid cells were dominant in the Ab+Pal resistant TME. We found that expression of several cytokines and chemokines involved in recruitment, differentiation and activation of myeloid cells (Fig. 2F, Csf1/M-CSF, Tgfβ2, Serpine2, Lgals3) were increased in response to Ab+Pal treatment. Studies have shown that tumor cell derived factors can promote bone marrow myeloid progenitor expansion and ultimately increase the number of circulating and tumor infiltrating immunosuppressive myeloid cells and contribute to disease progression ^14^. Additional studies of these cytokines/chemokines and other related immunomodulatory factors may provide greater insights into mechanisms of immunosuppressive myeloid cell accumulation during the tumor evolution/adaption and reveal potential targets for preventing disease progression or drug resistance. As clinical studies of anti-Her2 antibody and CDK4/6 inhibitor combination are still under way, the relevant data and samples of large patient cohorts are not yet available. Further investigation would be required to determine if immature myeloid cells infiltration is a general feature of disease progression/ therapeutic resistance as demonstrated in this study.

In summary, this study supports the necessity and provides potential value to use single-cell profiling to trace, characterize, and resolve tumor and TME evolution during the course of treatment, which could have a profound impact on future clinical decisions and rationally designed treatment strategies. Our preclinical findings indicated that targeting immature myeloid cells subverts immunosuppressive TME and restores the vulnerability of highly aggressive breast cancer to checkpoint blockade immunotherapy. Along with on-going clinical trial and patient tissue biopsy, we envision that similar prospective *in vivo* resistance modeling and rational regimen design informed by tumor and TME alterations, could facilitate future translational precision medicine for cancer patients.

## Materials and Methods

### Animal model and syngeneic tumor transplantation

FVB/N-MMTV-neu (202Mul) mice (Stock No: 002376) were purchased from Jackson Lab (Ben Harbor, ME). For tumor transplantation, treatment resistant tumors were excised from MMTV-neu mice and immediately cut into small pieces of 3-5 mm in diameter. Donor tumors were transplanted into 4^th^ mammary fat pad of MMTV-Neu mice (12 to14-week old). Incisions were closed with wound clips which were removed after 7-10 days. Mice were monitored daily for tumor establishment and then treatment was followed. Mouse experiments were performed in accordance with protocol approved by the University of Notre Dame IACUC committee.

### In vivo treatment

Anti-HER2/neu antibody (clone 7.16.4, BE0277), mouse IgG2a Isotype control (Catalog**#** BE0085), anti-CTLA 4 antibody (clone 9H10, BE0131), anti-PD-1 antibody (clone RMP1-14, BE0146), anti-Ly6G/Ly6C (Gr-1) antibody (clone RBC-8C5, BE0075), anti-CD3ε antibody (clone 145-2C11, BP0001) and polyclonal syrian hamster IgG (Catalog# BE0087) were purchased from BioXcell (West Lebanon, NH). Nulliparous female mice were enrolled for treatment when the spontaneous tumor reached a size of > 500 mm^3^. Anti-HER2/neu antibody or the isotype IgG control was intraperitoneally administered at 10 mg/kg body weight in PBS twice weekly. Palbociclib isethionate salt (LC laboratories, P-7766) was prepared in 50mM sodium lactate buffer and was given by oral gavage at a dose of 180 mg/kg every other day. Cabozantinib (LC laboratories, C-8901) dissolved in 30% (v/v) propylene glycol, 5% (v/v) Tween 80, and 65% (v/v) of a 5% (w/v) dextrose solution in water, was orally administered at daily dose of 30 mg/kg. For ICB treatment, Gr1^+^ cells and T-cell depletion experiments, anti-PD1, anti-CTLA 4, anti-Gr1 or anti-CD3ε antibodies (or their respective isotype IgG controls) were intraperitoneally administered at 200 μg per injection twice weekly, starting one day before anti-HER2/neu antibody and inhibitor treatment. The tumors were measured twice weekly using calipers. Tumor volume was calculated as length × width^2^ / 2. The volume of tumor when indicated treatment started was used as baseline for relative tumor volume calculation.

### Cell preparation

Cells for single cell RNA-seq were prepared by density centrifugation using Ficoll-Paque media (GE Healthcare, 17-5446-02) followed by magnetic-activated cell sorting (MACS) based separation or enrichment. In brief, fresh mammary tumors were resected and minced with sterile scissors into approximately 1- to 2- mm^3^ pieces, then enzymatically digested in DMEM/F12 medium (10 ml/g tumor) containing 5% FBS, 2 mg/ml collagenase (Sigma), 0.02 mg/ml hyaluronidase (Sigma), and 0.01 mg/ml DNase I (Sigma) for 30 minutes at 37°C with gentle agitation. Dissociated cells were centrifuged at 350 x *g* for 5 minutes with the brake on and discard supernatant. The pellet was re-suspended with 3-5 mL of pre-warmed TrypLE and incubated for 5 minutes. After adding 10 mL of DMEM/F12 medium supplemented with 2% FBS and passing through a 40 μm cell strainer (BD Biosciences), cells were centrifuged at 350 x *g* for 5 minutes and re-suspended in MACS buffer [phosphate-buffered saline (PBS) with 0.5% bovine serum albumin (BSA) and 2 mM EDTA]. Cell suspension was carefully layered on top of 15 ml Ficoll-Paque media solution in a 50-ml Falcon tube and centrifuged at 1000g for 10 minutes at room temperature with the break off. The buffy layer at the interface was transferred and washed with cold MACS buffer. Following Ficoll separation, dead cells were eliminated by using dead cell removal kit (Miltenyi Biotec, 130-090-101) per manufacturer’s instruction. The live cell fraction was then incubated with CD45 magnetically-labeled antibody (Miltenyi Biotec, 130-052-301) and passed through a LS magnetic column (Miltenyi Biotec, 130-042-401). The flow through fraction with enriched tumor cells (after depletion of CD45^+^ leukocytes) was collected. The cells retained in the column were then eluted as the isolated CD45^+^ tumor infiltrated leukocytes (TILs). For Gr1+ myeloid derived suppressor cell (MDSC) separation, the live cell fraction was subjected to a similar MACS based isolation by application of mouse MDSC isolation kit (Miltenyi Biotec, 130-094-538). Isolated cells were washed twice with cold MACS buffer and counted with a hemocytometer and diluted in cold PBS with 0.1% BSA and 2 mM EDTA at desired densities for Drop-seq.

### Drop-seq and sequencing analysis

Single-cell transcriptomic profiles were generated using Drop-seq protocol, as previously described ^51^. Briefly, enriched tumor cell suspensions (pooled from three or four tumors) as prepared above were loaded on the microfluidic device (fabricated in-house, CAD file from McCarroll Lab website: http://mccarrolllab.org/dropseq/) at approximately 100 cells/µL. CD45^+^ TILs and Gr1+ MDSCs were loaded at approximately 200 cells/µL (2 biological replicates for each treatment condition). Single cell suspension and uniquely barcoded microbeads (Chemgenes, MACOSKO201110) suspended in lysis buffer were co-encapsulated in droplets by the microfluidic device. The droplets serve as compartmentalizing chambers for RNA capture. Once droplet generation was complete, collected droplets were disrupted and RNA-hybridized beads were harvested. Reverse transcription was performed using Maxima H Minus Reverse Transcriptase (Thermo Fisher Scientific, EP0752) with template switching oligo (TSO). cDNA was amplified and PCR products were then purified using AMpure Beads (Beckman Coulter). After quantification on a BioAnalazyer High Sensitivity Chip (Agilent), samples were fragmented and amplified for sequencing with the Nextera XT DNA sample prep kit (Illumina). The libraries were purified, quantified, and then sequenced on the Illumina HiSeq 2500 or NextSeq 500. Sequencing format was 25-cycle read 1, 8-cycle index 1 and 50-cycle read 2. Base calling was done by Illumina Real Time Analysis (RTA) v1.18.64 and output of RTA was demultiplexed and converted to Fastq format with Illumina Bcl2fastq v1.8.4. Raw Drop-seq data were processed and aligned (STAR aligner) by following the standard Drop-seq pipeline (http://mccarrolllab.org/dropseq/). Briefly, reads were mapped to the mouse mm10 reference genome, then a digital gene expression data matrix was generated with counts of unique molecular identifiers (UMIs) for every detected gene (row) per cell barcode (column). We applied the knee plot method as recommended by the Drop-seq core computational protocol, which utilize the cumulative distribution of reads and identify an inflection point in the plot, to determine the number of cells (cell barcodes) represented in the expression matrix. Next, the Seurat R package ^52^ (V2.3.2, https://satijalab.org/seurat/) was used to perform data normalization, dimension reduction, clustering and differential expression analysis. Cells from corresponding treatment groups were merged into a single matrix. For tumor cells (sequenced by Illumina HiSeq 2500), genes with detected expression in at least 5 cells were included and cells with either less than 600 genes and 1500 UMI or more than 4000 genes and 20000 UMI were excluded. The percentage of reads aligned to mitochondrial genes per cells was calculated and cells with greater than 15% of transcripts derived from to mitochondrial genes ^53^ were filtered out. This resulted in 12638 genes across 4817 cells. Potential contaminating stromal cells were further removed based on the expression of *Pdgfra* (marker for fibroblast), *Pecam*/CD31(marker for endothelial cells), CD45 and CD11b (markers for leukocytes). We finally obtained 4711 cells for further analysis. For TILs in Fig. 2B (sequenced by Illumina HiSeq 2500), genes with detected expression in at least 2 cells were included and cells with either less than 400 genes and 1200 UMI or more than 4000 genes and 30000 UMI were excluded, and cells with greater than 10% of transcripts derived from to mitochondrial genes were removed. For Fig. 2G (sequenced by NextSeq 500), genes with detected expression in at least 2 cells were included, cells with either less than 500 genes and 1500 UMI or more than 5000 genes and 50000 UMI were excluded, and cells with mitochondrial genes greater than 10% were also removed. For Fig. 4B (sequenced by NextSeq 500), genes with detected expression in at least 10 cells were included, cells with either less than 400 genes or more than 5000 genes were excluded, and cells with mitochondrial genes greater than 10% were also removed. The filtered matrix was scaled to 10,000 molecules and log-normalized per cell to correct for the difference in sequencing depth between single cells.

### Gene set enrichment analysis

Single-sample gene set enrichment analysis (ssGSEA) ^54^ was run using GSVA v1.28.0 in R (https://www.bioconductor.org/packages/release/bioc/html/GSVA.html) using single-cell expression matrix with UMI values. We applied hallmark gene sets and canonical pathways from KEGG, REACTOME and BIOCARTA gene sets of the C2 collection of Molecular Signatures Database (MSigDB) (converted to mouse gene symbols) to each single cell to obtain enrichment score for each signature. Our custom and experimentally generated MDSCs signature was based on marker genes (top 300 differential expressed genes) of cell clusters by Seurat package. Drug target genes of FDA approved small molecular protein kinase inhibitors were adapted from The Blue Ridge Institute for Medical Research (http://www.brimr.org/PKI/PKIs.htm).

### CyTOF

Fresh or cryopreserved mammary tumors were enzymatically digested followed by density centrifugation and dead cell removal as aforementioned. Cells for CyTOF were washed and resuspended in Maxpar PBS (Fluidigm, 201058). Cells suspensions were incubated with Cell-ID Cisplatin (Fluidigm, 201064) for 5 minutes and then washed in Maxpar Cell Staining Buffer (Fluidigm, 201068). FC receptors were blocked by incubation with TruStain fcX in 100uL MaxPar Cell Staining Buffer for 15 minutes at room temperature. Cells were incubated with a cocktail of CyTOF antibodies (supplementary materials) for 30 min at room temperature and then washed in MaxPar Cell Staining Buffer. Optimal concentrations were determined for each antibody by titration. Cells were incubated with Cell-ID Cisplatin (Fluidigm, 201064) at 2.5 μM for 2.5 min for viability staining. Cells were resuspended and fixed in 1.6% PFA prepared in MaxPar PBS for 20 minutes and then Intercalator (Fluidigm, 201192B) dissolved in MaxPar Fix and Perm Buffer (Fluidigm, 201067) for 1 hour or overnight at 4 °C. Following nuclear labeling, cells were washed once in MaxPar Cell Staining Buffer and twice in MaxPar Water (Fluidigm, 201069). Samples were brought to 500,000 particulartes/mL in MilliQ water containing 0.1x EQ beads (Fluidigm, 201078) and run in 450μL injections on a CyTOF2 instrument. CyTOF data was analyzed and visualized using Cytobank Premium (Cytobank, Inc).

### Data and code availability

The single-cell RNA-seq data set has been deposited in the GEO data repository (accession number GSE122336). The custom scripts used for the described analysis are available from the corresponding authors upon reasonable request.

### Statistical analysis

Statistical tests were performed in GraphPad Prism version 7.0 or in R. Data were analyzed with two-tailed unpaired Student’s t tests when comparing means of two groups and one-way ANOVA when comparing more than two groups. Chi-square test was used to compare the proportion of cells. Survival curves were compared with the log-rank (Mantel-Cox) test. *P* values < 0.05 were considered significant.

## Supplementary Materials

### Materials and Methods

#### CyTOF antibodies

The following pre-conjugated antibodies purchased from Fluidigm were used in this study: CD45-089Y (3089005B, 30-F11); Ly-6G-141Pr (3141008B, 1A8,); CD11c-142Nd (3142003B, N418); CD45R-144Nd (3144011B, RA3-6B2); CD4-145Nd (3145002B, RM4-5); CD11b-148Nd (3148003B, M1/70); CD44-150Nd (3150018B, IM7); CD25-151Eu (3151007B, 3C7); CD3e-152Sm (3152004B, 145-2C11); PD-L1-153Eu (3153016B, 10F.9G2); CTLA-4-154Sm (3154008B, UC10-4B9); PD-1-159Tb (3159024B, 29F.1A12); Ly-6C-162Dy (3162014B, HK1.4); CX3CR1-164Dy (3164023B, SA011F11); NK1.1-165Ho (3165018B, PK136); c-Kit-166Er (3166004B, 2B8); CD8a-168Er (3168003B, 53-6.7); CD86-172Yb (3172016B, GL1); I-A/I-E-209Bi (3209006B, M5/114.15.2).

#### T-cell suppression assay

MDSCs were isolated from Ab+Pal treatment resistant tumors by Ficoll separation followed by using a mouse MDSC Isolation Kit (Miltenyi Biotec, 130-094-538). Isolate lymphocytes from spleen of wild-type FVB/N mice were labeled with carboxyfluorescein succinimidyl ester (CFSE) (2 μM) (Invitrogen, C34554), stimulated with CD3/CD28 magnetic beads (Invitrogen, 11452D), and cultured alone or with MDSCs at different ratios for 3 days. Cells were then collected and stained with anti-CD4-PE and anti-CD8a-APC (BioLegend, 100407 & 100711), CFSE intensity was quantified by flow cytometry and T-cell proliferation was analyzed.

#### Histology and immunohistochemistry

Tumor tissues were fixed in 10% neutral buffered formalin, processed routinely, and embedded in paraffin. H&E staining of paraffin-embedded tumor sections were used to quantify the hypocellular regions. Paraffin-embedded sections (4 μm) were subjected to antigen retrieval in a pressure cooker with sodium citrate buffer (PH=6.0) and incubated with antibodies specific for CD45 (BioLegend, 103103, 1:100), Ki-67 (DAKO, 1:200) overnight at 4°C. Biotin-conjugated secondary antibodies were used. Remaining steps were performed using Vectastain ABC kits (Vector Laboratories). Slides were counterstained with hematoxylin. To quantify the positive staining cells, the numbers of infiltrating CD3^+^ or Ki67^+^ cells were counted per field of view after examination of at least 10 fields of each section (200X), and the percentage of positive cells for Ki-67 was evaluated. Images were acquired using a Zeiss microscope with Axiovision software (Carl Zeiss, Inc.).

#### Immunofluorescence staining

FFPE sections were used for CD3 and CD8 IF staining. Sections (4 μm) were subjected to antigen retrieval in a pressure cooker with sodium citrate buffer (PH=6.0), blocked with R.T.U. Animal Free Blocker and Diluent (Vector Laboratories) for 1 h and then incubated with antibodies for CD3 (Abcam, ab16669, 1:300), CD8 (Thermo Scientific, 14-0808-82, 1:150) overnight at 4°C, followed by washing with PBS containing 0.05% TWEEN-20. Sections were then incubated with goat anti-Rabbit-Alexa Fluor 488 and goat anti-Rat-Alexa Fluor 594 (Thermo Scientific, A11034 and A11007, 1:500) in the blocking solution for 1 h at room temperature. After washing, sections were stained with DAPI to visualize nuclei. Cryosections were used for CD11b, Gr1, CD8 and granzyme B IF staining. For preparation of cryosections, dissected tissues were embedded in Tissue-tek O.C.T. (Electron Microscopy Sciences) and frozen on dry ice. Frozen tissues were stored at –80 °C until they were sectioned at 7 μm. For multicolored immunofluorescence staining, O.C.T. tumor cryosections were briefly air dried and fixed with 3% paraformaldehyde at room temperature for 15 min. Sections were then blocked with R.T.U. Animal Free Blocker and Diluent (Vector Laboratories) for 1 h and incubated with primary antibodies for CD11b-Alexa Fluor 488 (BioLegend, 101217, 1:50), Gr1-Alexa Fluor 594 (BioLegend, 108448, 1:50), CD8 (Thermo Scientific 14-0808-82, 1:150), granzyme B-FITC (BioLegend, 372205, 1:50). Secondary antibody goat anti-Rat-Alexa Fluor 594 (Thermo Scientific, A11007, 1:500) was used for CD8 staining. After washing, sections were stained with DAPI to visualize nuclei. Immunofluorescence imaging was performed on a multicolour fluorescent microscope (Leica DM5500 B). Five random fields were acquired from each biological sample for quantification of positive stained cells.

#### Flow cytometry sorting

Cryopreserved mammary tumors were enzymatically digested followed by density centrifugation. Cell suspensions were incubated with TruStain fcX (BioLegend, 101319) in 100uL MACS buffer (PBS with 0.5% BSA and 2 mM EDTA) for 15 minutes at room temperature. Cells were then incubated with pre-conjugated antibodies including CD45-APC (BioLegend, 103111), CD3-PE (Phycoerythrin) (BioLegend, 100205), F4/80-FITC (Fluorescein Isothiocyanate) (BioLegend, 123107), Gr1-APC/Cy (BioLegend, 108423). Flow cytometry sorting was performed on BD FACSAria. Single cells with CD3+, F4/80+ or Gr1+ (∼10,000) were sorted into tubes containing 50 uL lysis buffer in the PicoPure™ RNA Isolation Kit (Thermo Fisher Scientific, KIT0204).

#### Quantitative PCR

Total RNA was isolated using PicoPure™ RNA Isolation Kit (Thermo Fisher Scientific, KIT0204) and reverse-transcribed using qScript cDNA Synthesis Kit (Quantabio, 95047-100) following the manufacture’s protocol. Quantitative PCR was performed using 2x SYBR Green qPCR Master Mix (Bimake, B21202). Primers were synthesized by Integrated DNA Technologies and the sequences are listed in Supplementary Table 2. Gene expression level was calculated relative to β-actin using ΔCt values.

## Author contributions and Acknowledgments

QW and SZ conceived the original hypothesis and designed experiments. QW, IHG, SMG, JL, LS, and JH performed experiments. QW and SZ analyzed data. XL provided direction and guidance to this study. QW and SZ wrote and revised the manuscript. SZ supervised the study. We would like to thank Zhang Lab members for scientific insights and support. For insightful technical assistance, we thank Jacqueline Lopez, M.S., Samuel W. Brady, Ph.D., Andrea Gunawan, M.S. and Charles R. Tessier, Ph.D. We are additionally grateful for the use of the following core facilities: Notre Dame Genomics and Bioinformatics Core Facility, Notre Dame Freimann Life Sciences Center, Harper Cancer Research Institute Biorepository, Indiana University Simon Cancer Center Core Facility, Michigan State University Genomics Core. This work was partially funded by NIH R01 CA194697-01 (SZ), NIH R01 CA222405 - 01A1(SZ), Notre Dame CRND Catalyst Award (SZ and IHG), NIH CTSI core facility pilot grants (SZ), U54 pilot grant U54CA209978 (SZ), Notre Dame ADT grant (SZ), NIH CTSI Postdoc Challenge award (QW). We would additionally like to acknowledge and thank the Dee Family endowment (SZ).

---

**Fig. S1.**
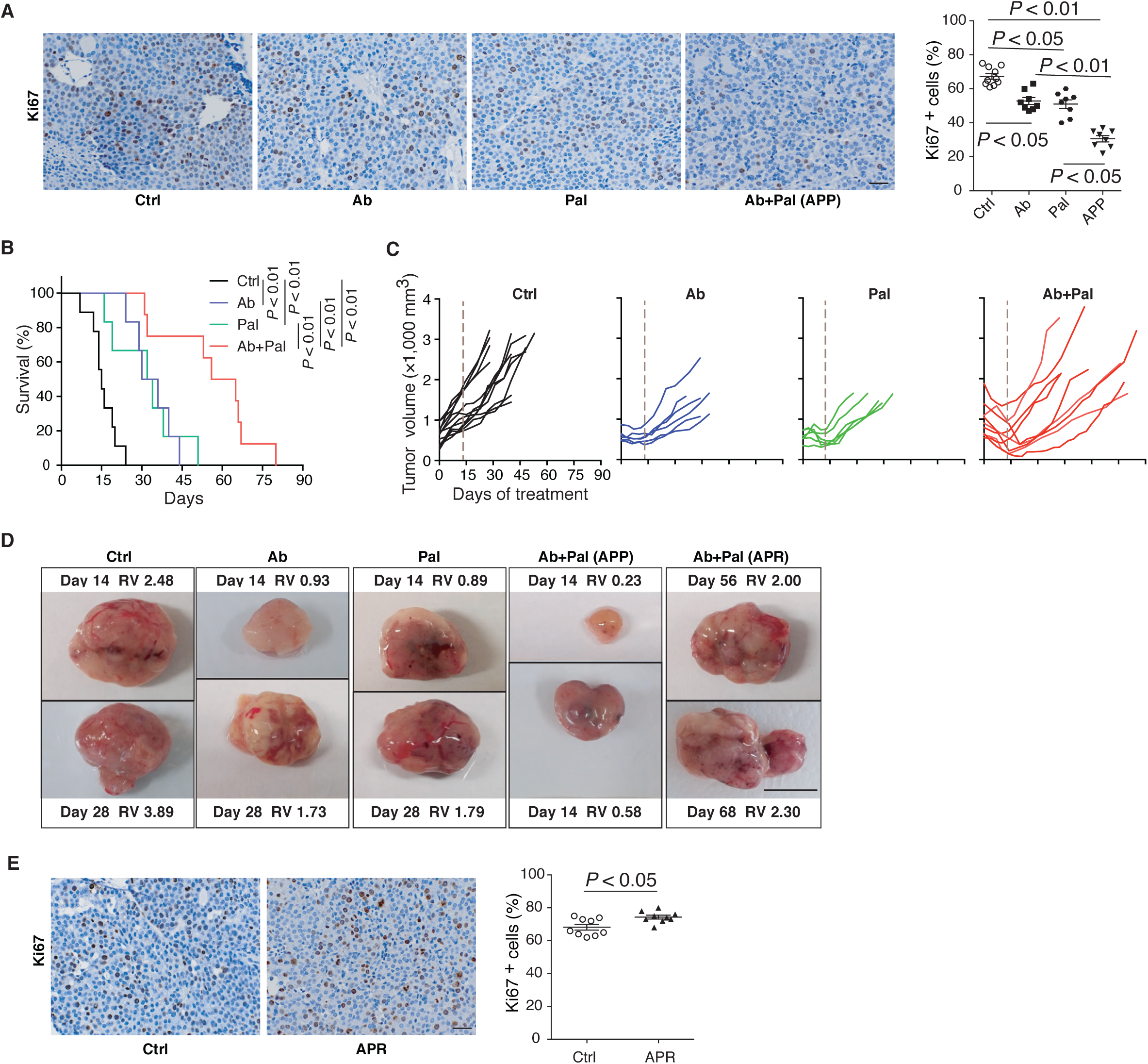
Response and resistance to anti-Her2/Neu antibody plus CDK4/6 inhibitor Palbociclib combination in spontaneous Her2/Neu-positive breast cancer. (**A**) Representative images and quantification of Ki67 immunohistochemistry staining. MMTV-Neu mice (bearing tumor > 500 mm3) were treated with Ctrl, Ab or Pal alone, or Ab+Pal for 14 days, then tumors were harvested. Ctrl, vehicle treated control; Ab, anti-Her2/Neu antibody; Pal, CDK4/6 inhibitor Palbociclib. Scale bar, 20 μm. *P*-value by one-way ANOVA with Tukey’s test. (**B**) Survival time to doubled tumor volume (n=9,6,6 and 8 for Ctrl, Ab, Pal and Ab+Pal). *P*-value by log-rank (Mantel-Cox) test. (**C**) Individual tumor growth kinetics with Ctrl (n=12), Ab (n=6) or Pal (n=5) alone, or Ab+Pal (n=10) treatment. Dash line indicating 14-day’s treatment. (**D**) Representative images of tumors with indicated treatment. RV: relative tumor volume. RV = tumor volume at the indicated time (days of treatment) / volume at the start point of treatment. Scale bar, 1cm. (**E**) Representative images and quantification of Ki67 immunohistochemistry staining on tumors derived from mice with 7 weeks’ treatment. Scale bar, 20 μm. *P*-value by Student’s t-test.

**Fig. S2.**
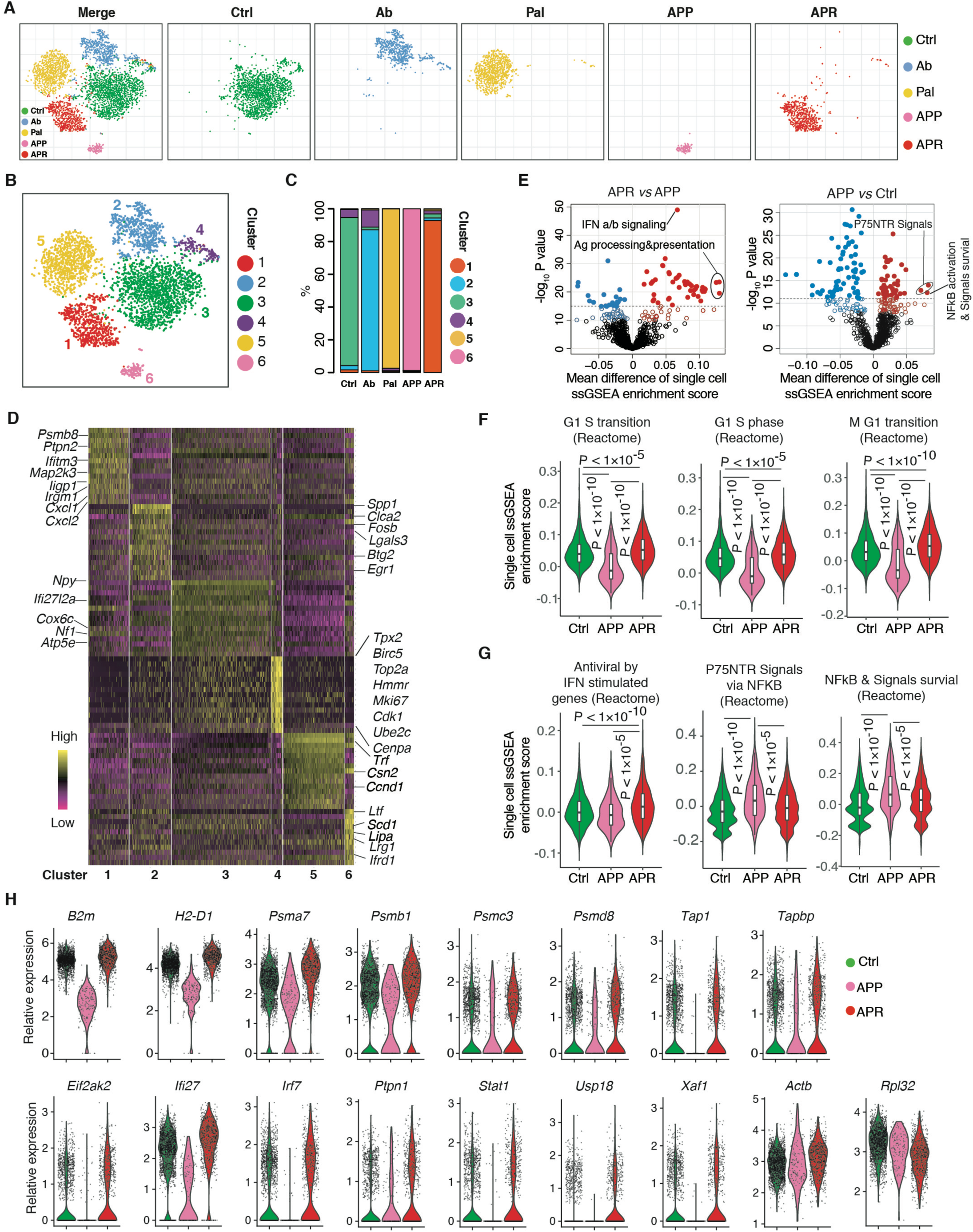
Single-cell RNA sequencing of tumor cells from responsive and resistant tumors to anti-Her2/Neu antibody plus CDK4/6 inhibitor Palbociclib combination treatment. (**A**) Clustering of 4711 tumor cells by single-cell expression profiles and I-distributed stochastic neighbor embedding (t-SNE) plots colored by different treatment conditions [merged (left) or subdivided into individual treatment group (right)] are shown. Each point represents a single cell. (**B**) t-SNE plot from (A) is colored by clusters.(**C**) Abundance of each cluster [as clustered in (B)] in tumors with indicated treatment.(**D**) Heatmap of top differentially expressed (marker) genes of each cluster. Each column represents a single cell.**(E**) Volcano plots comparing ssGSEA enrichment score for 1053 canonical pathways/gene sets of the C2 collection of Molecular Signatures Database between APR and APP (left), APP and Ctrl (right), based on single cell RNA-Seq data. Each point represents one pathway/gene set. X-axis, mean difference of single cell ssGSEA enrichment score; Y-axis, −log1O (P-value by t-test).(**F and G**) ssGSEA enrichment score violin plots for single cells from each treatment group for cell cycle related signatures (F) and indicated signatures (G). P-value by Student’s I-test (two-tailed).(H) Expression of genes involved in antigen processing and presentation (up panel), IFN signaling and response (low panel) in individual cells from tumors with indicated treatment. Each point represents a single cell. Expression of house keep genes Actb and Rpl32 in individual cells are also shown.

**Fig. S3.**
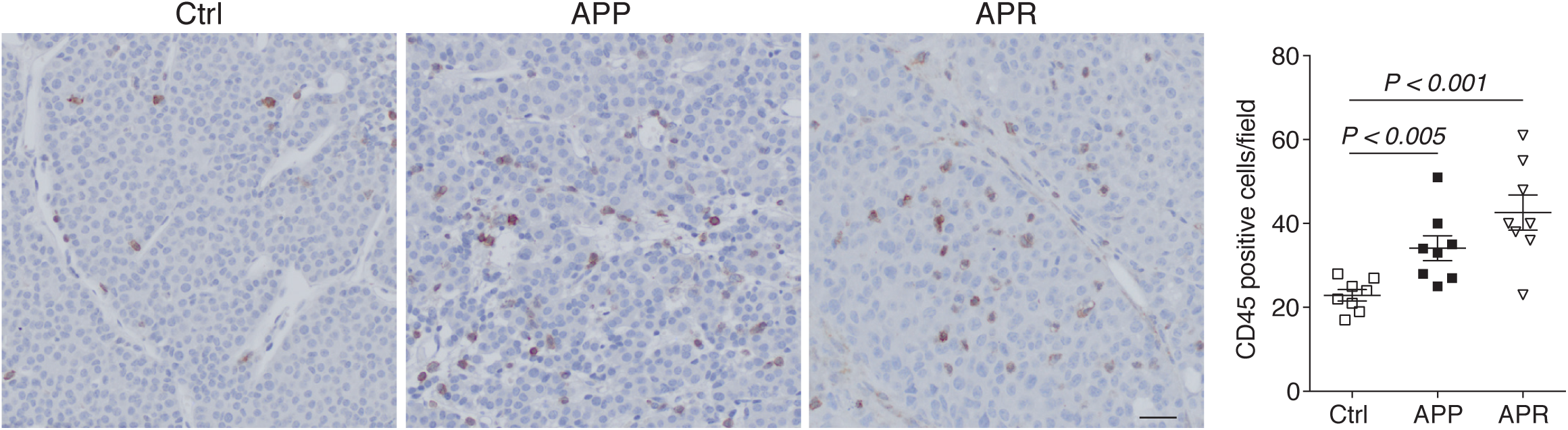
Ab+Pal treatment increased tumor infitratiing leukocytes. Representative images and quantification of CD45 immunohistochemistry staining for Ctrl, APP and APR tumors. Scale bar, 20 μm. *P*-value by one-way ANOVA with Tukey’s test.

**Fig. S4.**
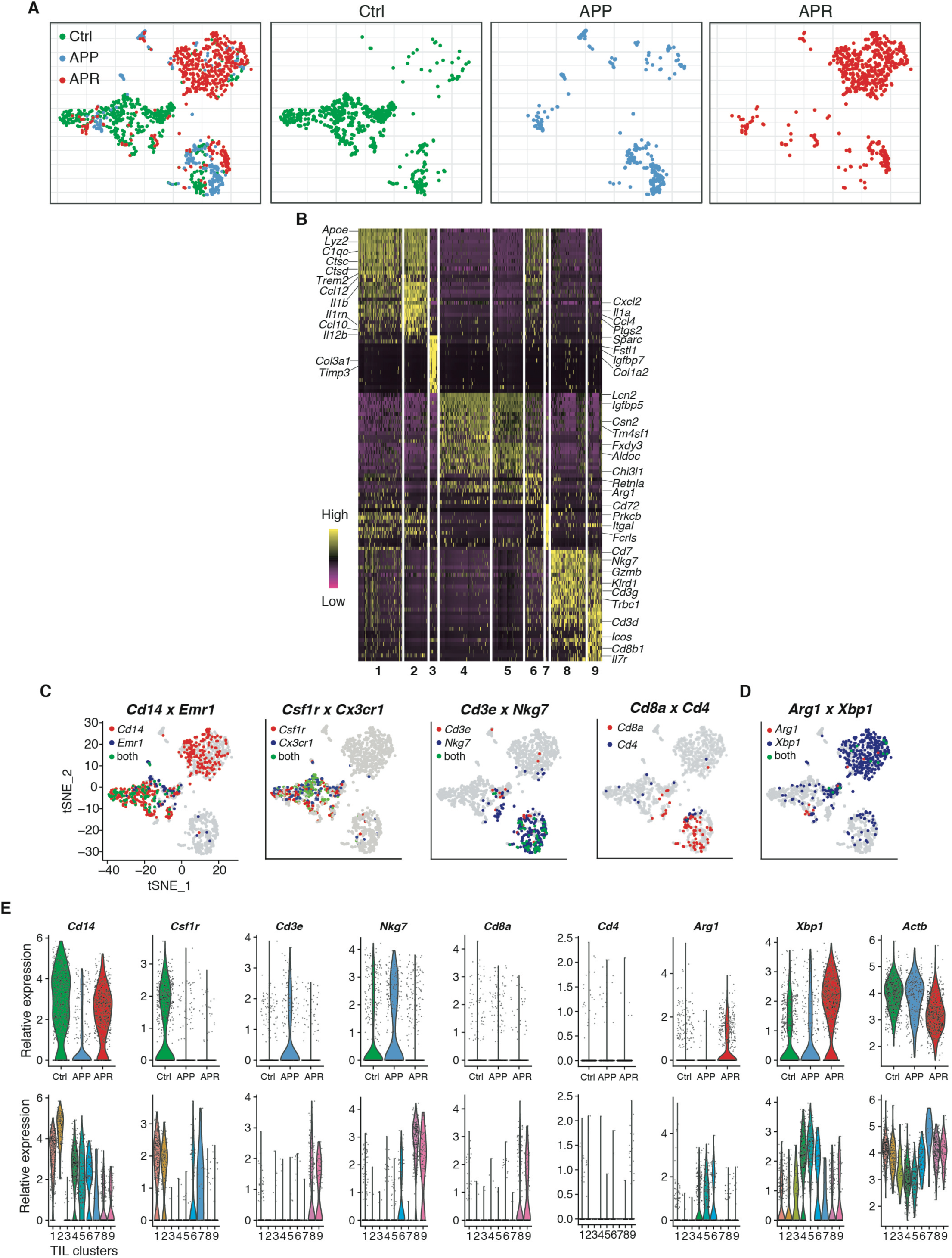
Single-cell RNA sequencing of tumor infiltrated leukocytes derived from Ab+Pal treatment responsive and resistant tumors. (**A**) Clustering of 1444 Tlls derived from Ctrl, APP and APR tumors and t-SNE plots colored by different phenotypes [merged (left) or subdivided into individual group (right)] are shown]. Each point represents a single cell. (**B**) Heatmap of top differentially expressed (marker) genes of each cluster from experiment (A). Each column represents a single cell. **(C** and **D)** Expression of key marker genes used for immune cell-type identification and annotation was overlaid on t-SNE plots.(E) Expression distribution of key marker genes [as mentioned in (C and D)] and house keeping gene Actb among tumors with different phenotypes (up) and different TIL-clusters were shown. Each dot represents a single cell.

**Fig. S5.**
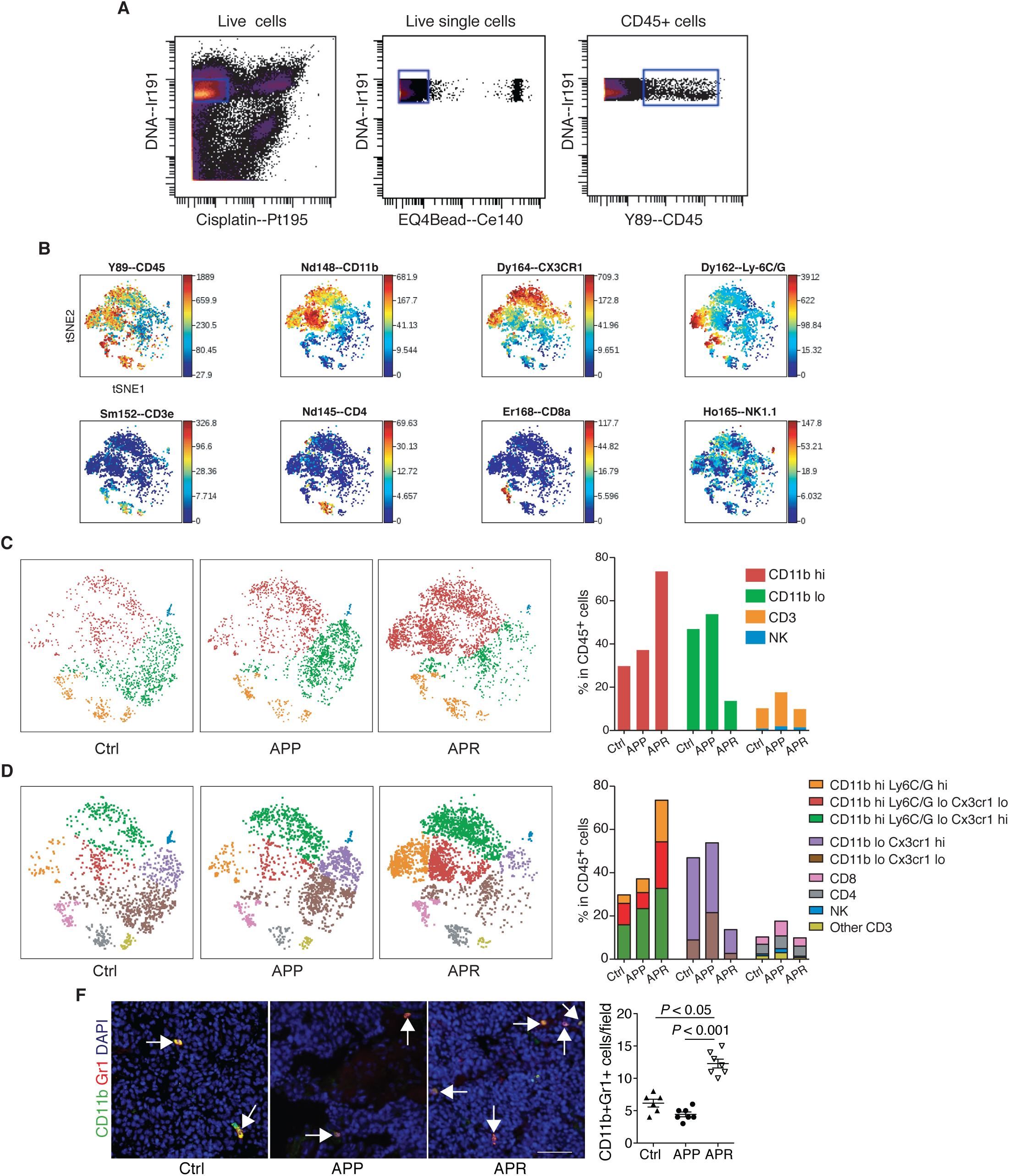
Characterization of infiltrating immune cell populations in responsive and resistant tumors to Ab+Pal combination treatment. (**A**) Preliminary gating for mass cytometry samples. Iridium 191 and cisplatin 195 were used to select live single cells and subsequently CD45+ cells were selected for further analysis. (**B**) Representative plots showing the expression of markers used for cell characterization/gating overlayed on viSNE plots. viSNE plots were generated by Cytobank, based on the t-SNE algorithm. Each point represents a single cell and the color gradient represents marker’s intensity (expression level). (**C**,**D**) Classification of CD45+ tumor infiltrated immune cell populations on viSNE plots and quantification. (**F**) Representative images and quantification of CD11b and Gr1 immunofluorescence staining for Ctrl, APP and APR tumors. Arrows indicate CD11b and Gr1 double positive cells. Scale bar, 50 μm. P-value by one-way ANOVA with Tukey’s test.

**Fig. S6.**
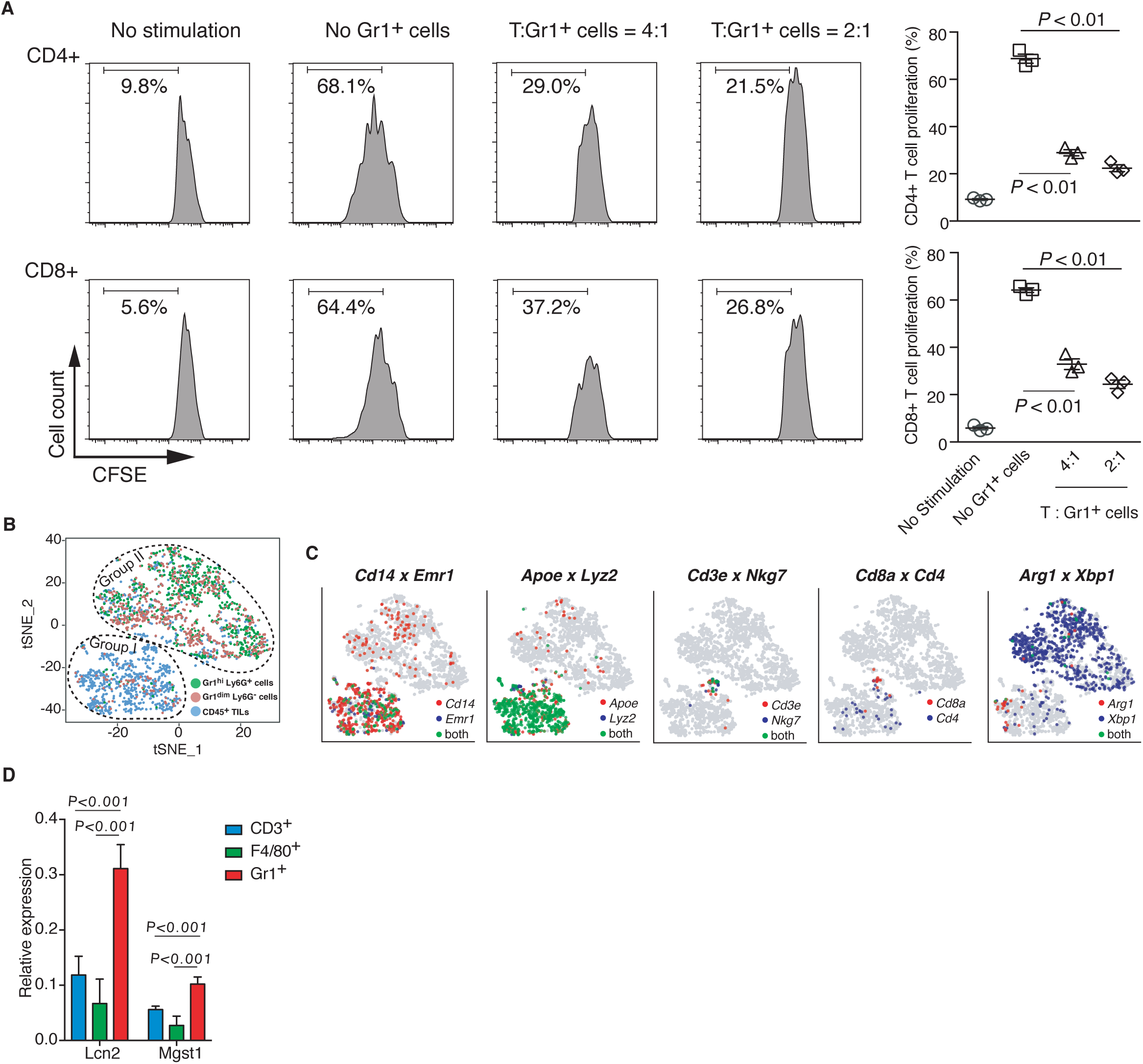
Functional characterization and single-cell RNA sequencing of tumor infiltrated MDSCs. (**A**) Gr1+ MDSCs isolated from Ab+Pal treatment resistant tumors inhibited CD3/CD28 stimulated proliferation of CD4+ and CD8+ T cells in vitro. Left, representative flow cytometry histograms measuring carboxyfluorescein succinimidyl ester (CFSE); right, quantification of T-cell proliferation. *P*-value by Student’s t-test. (**B**) t-SNE plot of single cell RNA sequencing data from CD45+ TILs, Gr1high Ly6G+cells and Gr1dim Ly6G-cells. Each point represents a single cell. (**C**) Expression of key marker genes used for immune cell-type identification and annotation was overlaid on t-SNE plots as shown in (B). (**D**) mRNA levels of Lcn2 and Mgst1 in sorted T cells (CD3+), macrophages (F4/80+) and MDSCs/IMCs (Gr1+) from APR tumors were quantified by RT-PCR. *P*-value by one-way ANOVA with Tukey’s test.

**Fig. S7.**
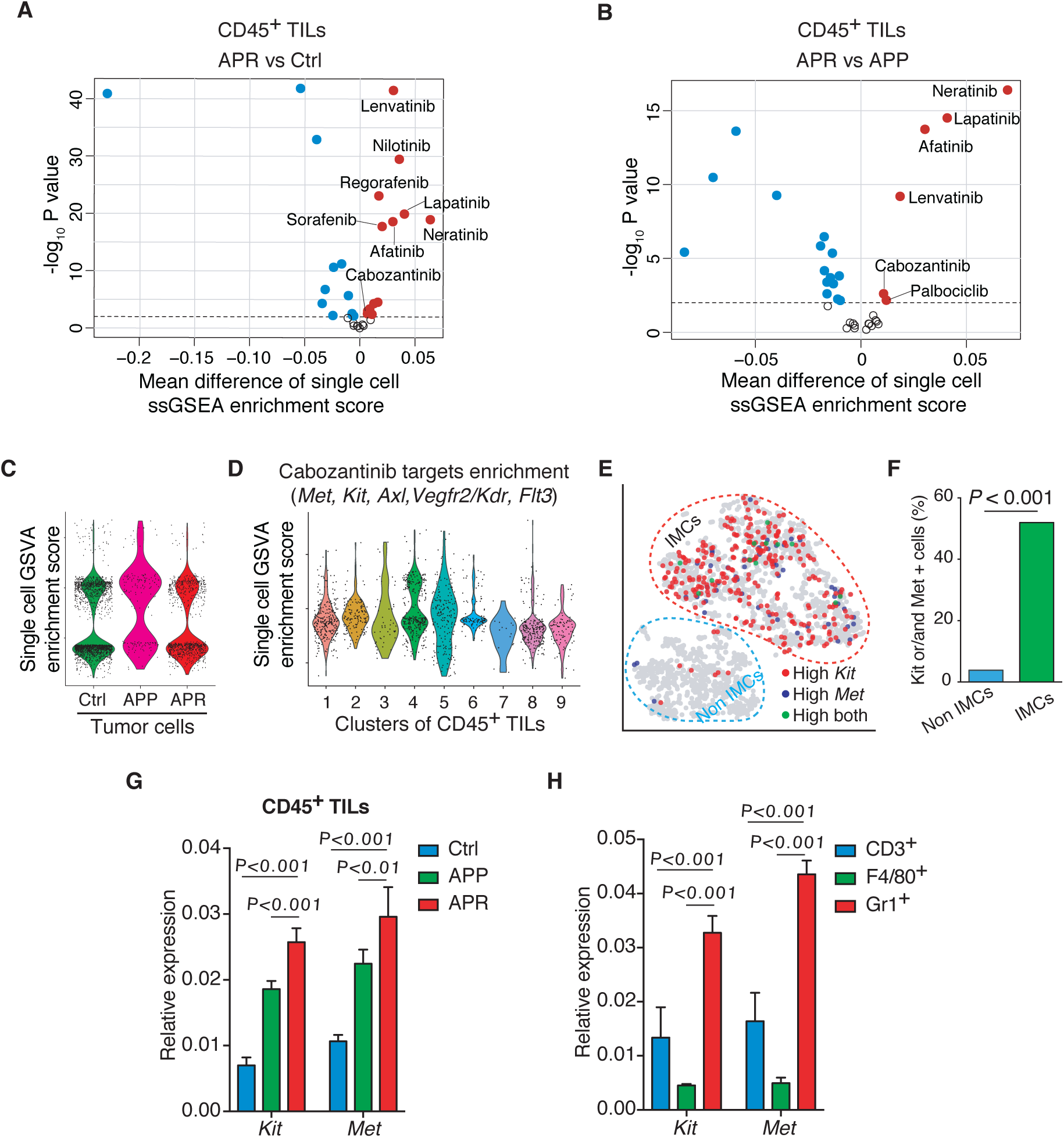
Identification of cabozantinib as a potential IMCs-targeted drug by single-cell RNA-seq analysis. (**A,B**) Volcano plots comparing enrichment score for drug target genes of FDA approved protein kinase inhibitors between TILs from resistant and control tumors (A, APR vs Ctrl), and from resistant and responsive tumors (B, APR vs APP). Each point represents an inhibitor. X-axis, mean difference of ssGSEA enrichment score between groups; Y-axis, −log10 (P-value by t-test). (**C**) Enrichment analysis of cabozantinib target genes across tumor cells grouped and plotted by different phenotypes. Each point represents a single cell. (**D**) Distribution of enrichment score for cabozantinib target genes among TIL-clusters as identified in Fig.2A. (**E**) Distribution of Kit and/or Met expression cells on t-SNE of single cell RNA sequencing data from CD45+ TILs and Gr1+ cells (experiment of Fig.2G). Each point represents a single cell. **F)** Abundance of Kit and/or Met expressing cells in IMCs and non-IMCs as annotated in (E). *P*-value by Chi-square test. (**G, H**) mRNA levels of Kit and Met in CD45+ TILs form Ctrl, APP and APR tumors (**G**) and in sorted T cells (CD3+), macrophages (F4/80+) and MDSCs/IMCs (Gr+) from APR tumors (**H**) were quantified by RT-PCR. *P*-value by one-way ANOVA with Tukey’s test.

**Fig. S8.**
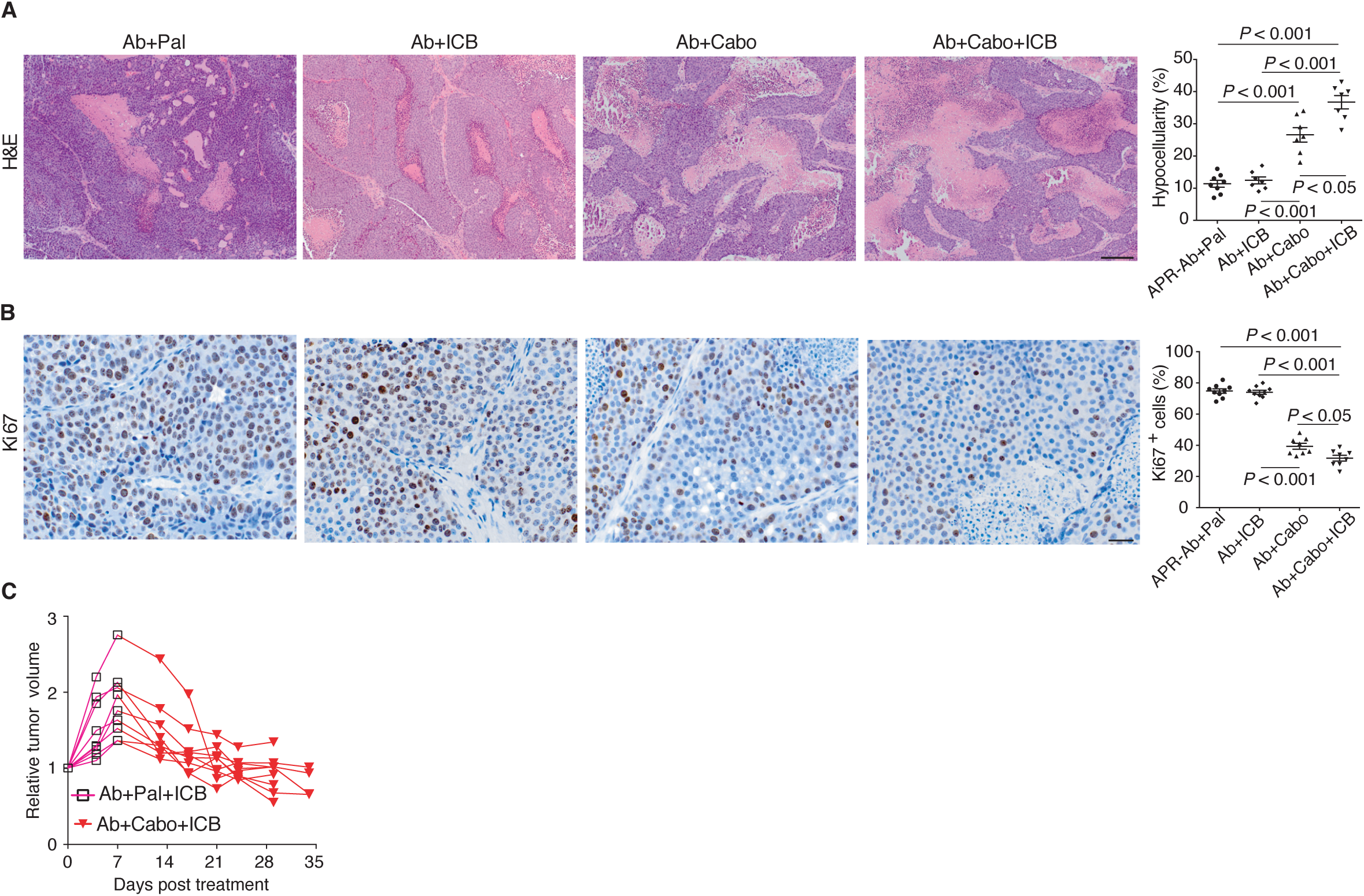
Therapeutic activity of Ab+Cabo or Ab+Cabo+ICB against Ab+Pal resistant tumors. (**A**) Representative images of H&E staining for tumors with indicated treatment and quantification of hypocellularity. Scale bar, 100 μm. (**B**) Representative images and quantification of Ki67 immunohistochemistry staining tumors with indicated treatment. Scale bar, 20 μm. (**C**) Relative volumes of Ab+Pal resistant tumors after sequential treatment with Ab+Pal+ICB and Ab+Cabo+ICB (as in Fig.3F). Ab+Pal resistant tumors were first treated with Ab+Pal+ICB for 1 week then switched to Ab+Cabo+ICB treatment for 3 weeks. Cabo, protein kinase inhibitor cabozantinib; ICB, immune checkpoint blockades, cocktail of anti-CTLA4 and anti-PD-1 antibodies. *P*-value by one-way ANOVA with Tukey’s test.

**Fig. S9.**
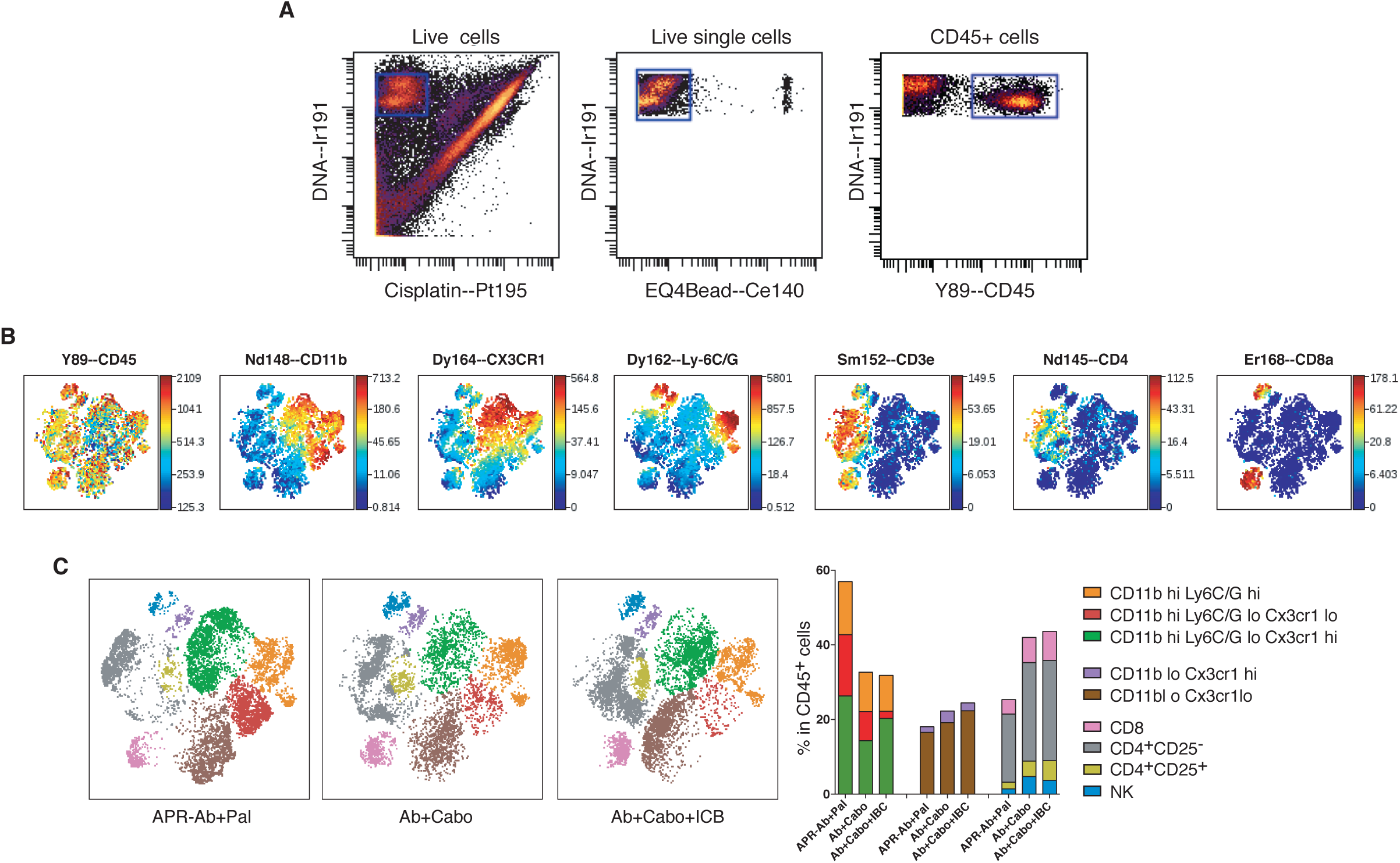
Characterization of infiltrating immune cell populations in tumors after Cabo and ICB treatment. (**A**) Preliminary gating for mass cytometry samples after Cabo and ICB treatment. Iridium 191 and cisplatin 195 were used to select live single cells and subsequently CD45+ cells were selected for further analysis. (**B**) Representative plots showing the expression of markers used for cell characterization/-gating overlayed on viSNE plots. viSNE plots were generated by Cytobank, based on the t-SNE algorithm. Each point represents a single cell and the color gradient represents marker’s intensity (expression level). (**C**) Classification of CD45+ tumor infiltrated immune cell populations on viSNE plots (left) and quantification (right).

**Fig. S10.**
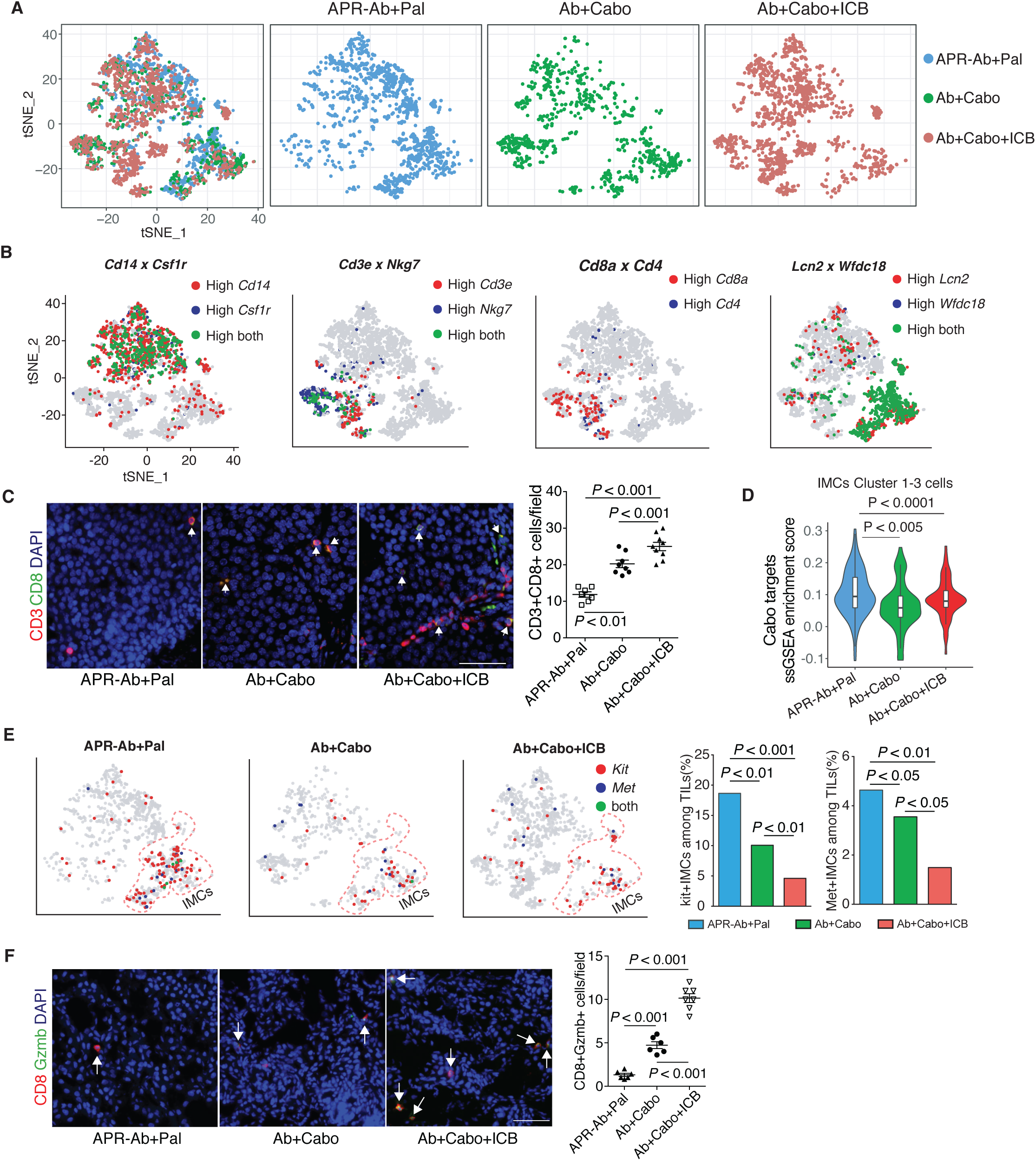
Single cell RNA sequencing of tumor infiltrated leukocytes after Cabo and ICB treatment. (**A**) t-SNE plots of 3168 TILs derived from resistant tumors with continuous Ab+Pal, Ab+Cabo or Ab+Cabo+ICB treatment [merged (left) or subdivided into individual group (right)] were shown]. Each point represents a single cell. (**B**) Expression of key marker genes used for immune cell-type identification and annotation was overlaid on t-SNE plots. (**C**) Representative images and quantification of CD3 and CD8 immunofluorescence staining. Arrows indicate CD3 and CD8 double positive cells. Scale bar, 50 μm. *P-*value by one-way ANOVA with Tukey’s test. (**D**) Enrichment scores of cabozantinib target genes among tumor infiltrated IMCs after indicated treatment. The P values showed were determined by Student’s t-test. (**E**) Distribution of Kit and/or Met expression cells on t-SNE plot of single cell RNA sequencing data from (A), and abundance of Kit or Met expressing IMCs within CD45+ TILs after indicated treatment. *P*-value by three-sample Chi-square test. (**F**) Representative images and quantification of CD8 and Granzyme B immunofluorescence staining. Arrows indicate CD8 and Granzyme B double positive cells. Scale bar, 50 μm. *P-*value by one-way ANOVA with Tukey’s test.

**Supplementary Table 1.**
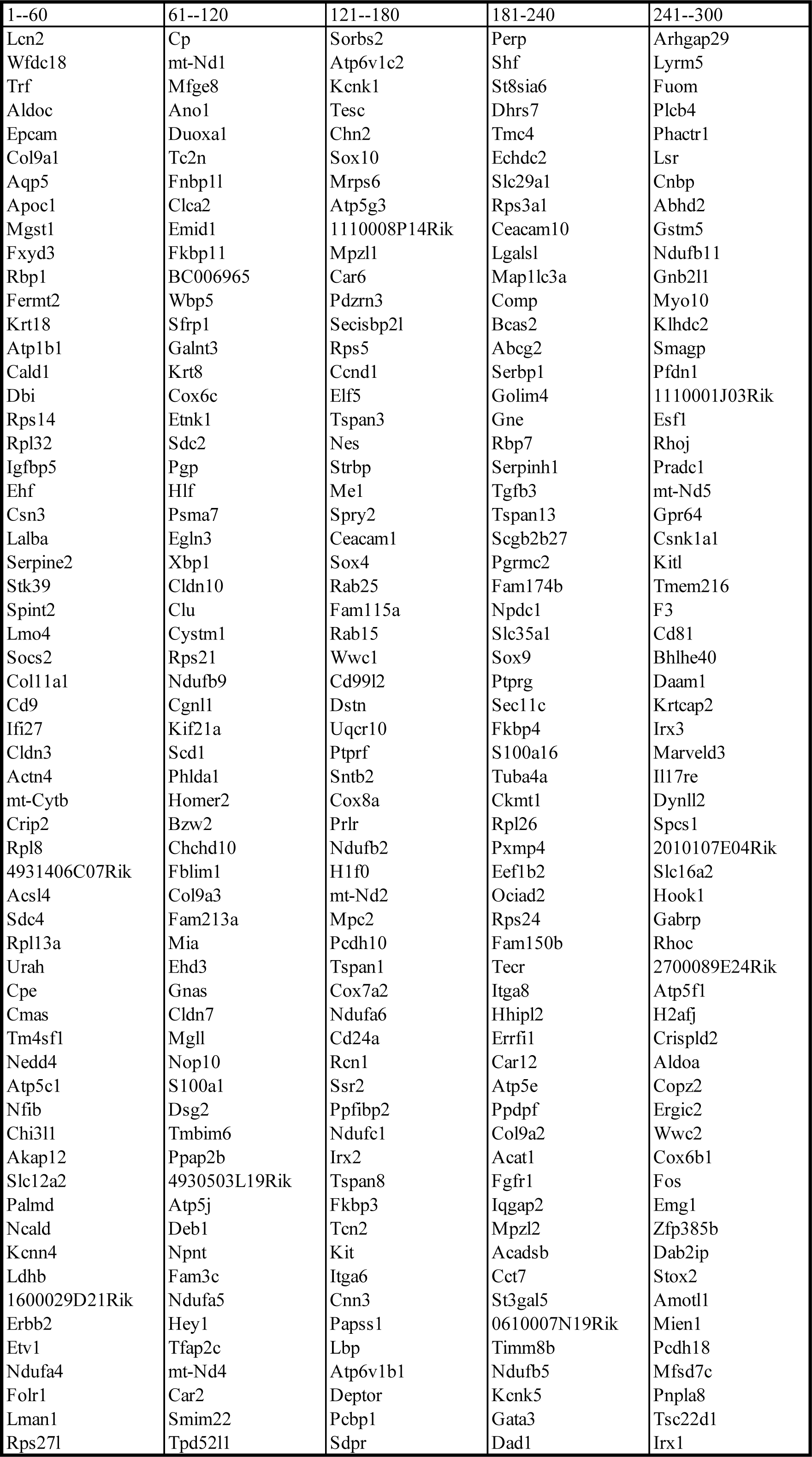
Gr-1+ MDSCs signature (Top 300 differentially expressed genes of MDSCs clusters by scRNA-seq as shown in Fig. 2H)

**Supplementary Table 2.**
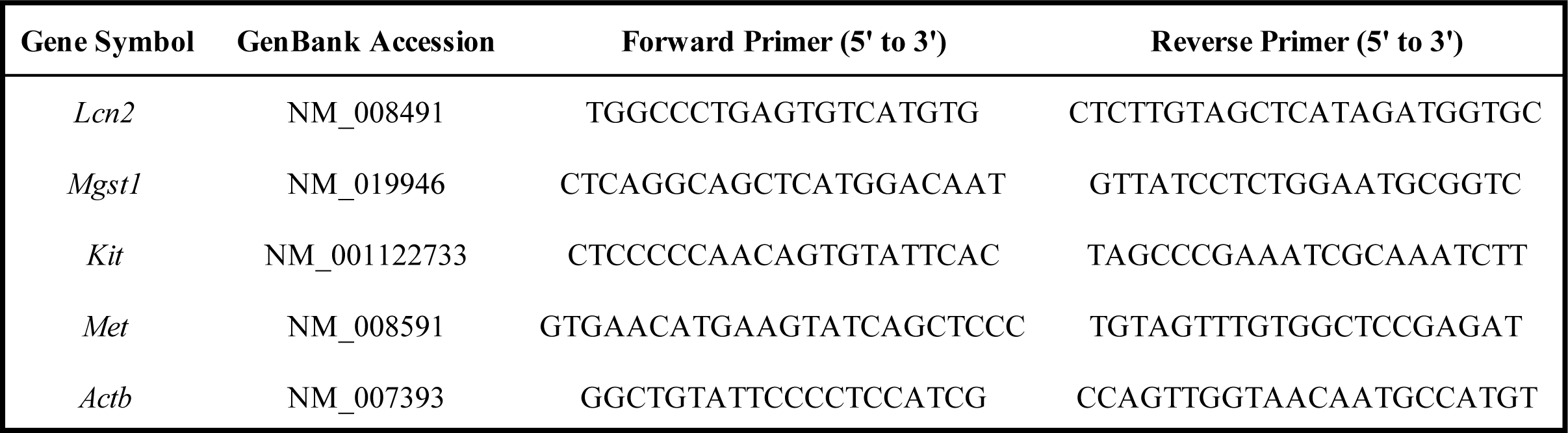
Primer sequences

